# Mycobacterial MutT1 mediated dephosphorylation of the sensor kinases reveals a new link in the regulation of the two-component signaling in bacteria

**DOI:** 10.1101/2025.03.06.641775

**Authors:** Elhassan Ali Fathi Emam, Devendra Pratap Singh, Deepak K. Saini, Umesh Varshney

**Affiliations:** Departments of Microbiology and Cell Biology, Bangalore, 560012, India; Developmental Biology and Genetics, Bangalore, 560012, India; Centre for Biosystems Science and Engineering, Indian Institute of Science, Bangalore, 560012, India; Jawaharlal Nehru Centre for Advanced Scientific Research, Bangalore, 560064, India

**Keywords:** *Mtb*MutT1, *Msm*MutT1, RHG histidine phosphatase, sensor kinase, response regulator, SenX3-RegX3, PhoPR

## Abstract

Bacterial pathogens such as *Mycobacterium tuberculosis* majorly rely on two-component signaling (TCS) systems to sense and generate adaptive responses to the dynamic and stressful host environment. TCS comprises a sensor histidine kinase (SK) that perceives the environmental signal, and a response regulator (RR) that modulates the target gene expression. TCS occurs via a phosphotransfer event from SK to RR. However, the mechanisms that regulate phosphotransfer events are not well understood. We explored the role of MutT1, originally characterized to hydrolyze oxidized GTP (8-oxo-GTP) and dGTP (8-oxo-dGTP), in TCS regulation. Unlike other MutT proteins, mycobacterial MutT1 comprises two domains (N-terminal domain, NTD; and C-terminal domain, CTD). Structurally, MutT1 NTD is like MutT proteins in other organisms. However, the MutT1 CTD is similar to *E. coli* SixA, a histidine phosphatase with an RHG motif. We show that MutT1 CTD dephosphorylates many SKs and impacts expression of their target genes, highlighting the role of MutT1 in regulating TCS. These novel findings are of special significance because they provide us with an extrinsic phosphatase mechanism to reset TCS signaling. The study reveals an intricate interplay between an enzyme that sanitizes the cellular nucleotide pool, and bacterial signaling pathways, offering insights into the adaptation mechanisms.

## Introduction

*Mycobacterium tuberculosis*, a pathogenic bacterium, possesses exceptional survival strategies to evade host defenses. It consistently monitors the host environment to orchestrate expression of genes required for its survival. The two-component signaling (TCS) systems make a major contribution to acquiring this adaptability by regulating transcriptional networks [1–4]. TCSs comprise a sensor histidine kinase (SK) that perceives signal, and a response regulator (RR) that modulates target gene expression. SKs are often a transmembrane protein, containing a variable sensor domain and a conserved kinase domain, while RRs are cytosolic proteins consisting of a conserved receiver domain and variable effector domain [3, 5]. Upon perceiving signal through its sensory domain, the kinase domain of the SK autophosphorylates itself at a conserved histidine (His) residue. This is then followed by phosphotransfer event wherein the kinase domain of SK transfers the same phosphate group to a conserved aspartate (Asp) residue of its partner RR. The phosphorylation of RR alters its DNA binding properties, leading to modulation of target gene expression that facilitates an adaptive response to the external stimuli [3, 5–8]. In addition, the positive autoregulation of the TCS itself is a key aspect of the adaptive response. The SK and RR genes of a TCS pathway are generally present in an operonic organization. Interestingly, the phosphorylated RR, acting as a transcription factor, positively autoregulates its own operonic expression, thereby enhancing the TCS sensitivity and the adaptive response to the external stimuli [9]While a positive feedback in TCSs is advantageous under strong and persistent stimuli, it can produce disproportionate response to a weak or fleeting signal and impact bacterium fitness [9–14]. This raises an important question of how bacteria may prevent such disproportionate responses. Are there physiological mechanisms that regulate the cellular levels of phosphorylated SKs (SKs-P), which in turn may downregulate its activity in transfer of the phosphate group to the downstream partner RRs to moderate gene expression.

*M. tuberculosis* genome encodes for 12 paired TCSs, 2 orphaned SKs and 4 orphaned RRs [15]. The TCS such as HK1-HK2-TcrA, KdpED, and TrcRS, and the orphaned SKs and RRs, lack comprehensive characterization [4]. However, other TCSs, namely PdtaRS, DosRST, MprAB, TcrXY, NarLS, PrrAB, MtrAB, PhoPR and SenX3-RegX3 have been better characterized for their roles in virulence [4, 6, 16–28]. TCSs exhibit significant sequence and structural similarities in their conserved domains, are known for the crosstalk among SK and RR proteins, and recognised to offer possible ways of signal integration or diversification in bacteria [4, 8, 29–32].

Mycobacterial MutT1 comprises two distinct domains: the N-terminal domain (NTD) having the Nudix hydrolase motif (GX_5_EX_7_REUX EEXGU, where U represents a hydrophobic amino acid and, X is any amino acid) and the C-terminal domain (CTD) containing the RHG histidine phosphatase motif [33] (Fig. 1A). MutT1 NTD hydrolyses 8-oxo-(d)GTP (8-oxo-GTP or 8-oxo-dGTP) to prevent 8-oxo-G incorporation in nucleic acids [33, 34]. Recently, we identified a novel role of MutT1 NTD in regulating the activity of nucleoside diphosphate kinase (NDK) by dephosphorylating NDK-P. Further, we showed that the MutT1 CTD contributes structurally to regulating this activity [35]. Despite these insights, the primary role of MutT1 CTD remains unknown. MutT1 CTD revealed an extraordinary structural resemblance to *E. coli* SixA (*Eco*SixA) [33]. Both the MutT1 CTD and SixA feature an RHG histidine phosphatase motif (Fig. 1B). *Eco*SixA exhibits phosphatase activity towards ArcB, a SK in ArcAB TCS [36]. To date, there are no reports of TCSs regulation by the RHG based histidine phosphatase activity of the MutT1 in mycobacteria.

**Fig. 1:**
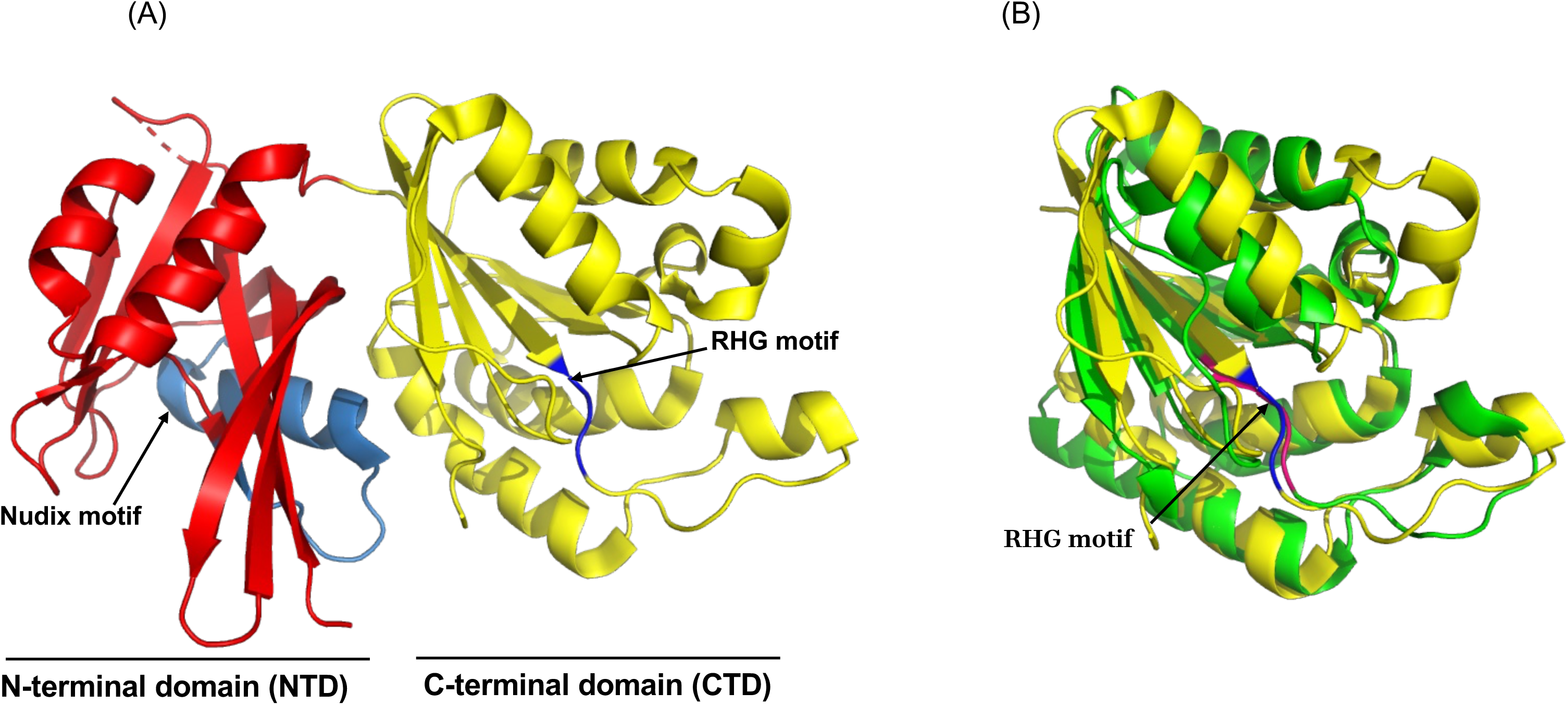
Structural comparison between mycobacterial MutT1 and *E. coli* SixA proteins. (A) A representation of crystal structure of *M. smegmatis* MutT1 (PDB ID-5GG5) [33]. The *Msm*MutT1 NTD (red) possesses the Nudix hydrolase motif highlighted in light blue. The *Msm*MutT1 CTD represented in yellow contains RHG motif (blue). (B) Structural comparison of *Msm*MutT1 CTD (yellow, PDB ID-5GG5 [33]) and *Eco*SixA (green, PDB ID-1UJB) [39]. The RHG motifs are highlighted in blue (*Msm*MutT1 CTD) and red (*Eco*SixA).

In this study, we aimed to determine if mycobacterial SKs serve as suitable substrates for MutT1. We show that MutT1 CTD indeed possesses phosphatase activity against a subset of SKs. Further, we show a significant role of MutT1 in modulating the activity of mycobacterial TCSs.

## Results

### Mycobacterial MutT1 dephosphorylates a group of SKs

Based on the structural similarity of mycobacterial MutT1 CTD with *Eco*SixA, and the presence of RHG motif [33], we aimed to investigate if MutT1 participates in regulation of mycobacterial TCSs. We screened eleven mycobacterial SKs as potential substrates for MutT1 (Figs. 2 and S1). As in the earlier studies [7, 30], we have used only the kinase domains of the mycobacterial SKs (except for PdtaS for which full length SK was used). We interrogated histidine phosphatase activity of MutT1 CTD using autophosphorylated SKs with both *M. tuberculosis* MutT1 (*Mtb*MutT1) and its ortholog from *M. smegmatis* (*Msm*MutT1). Interestingly, both *Mtb*MutT1 and *Msm*MutT1 decreased the phosphorylation levels of autophosphorylated *Mtb*SenX3 (*Mtb*SenX3-*P*) and *Mtb*PhoR (*Mtb*PhoR-*P*) (Figs. 2A and 2B). In contrast, when autophosphorylated *Mtb*NarS (*Mtb*NarS-*P*) and *Mtb*PdtaS (*Mtb*PdtaS-*P*) were used, no changes in their phosphorylation levels were detected (Figs. 2C and 2D). Further, MutT1 displayed phosphatase activity on other SKs such as *Mtb*MtrB, MprB, DosS, and PrrB (Figs. S1A, S1B, S1C, S1D and S1E) but not on *Mtb*DosT, TcrY, and KdpD (Figs. S1F, S1J and S1K). These observations suggest specificity of MutT1-mediated release of *Pi* from the autophosphorylated SKs (Figs. 3A and 3B, compare lanes 3 with lanes 1 and 2). And, in these reactions, phosphorylation of MutT1 itself was not detected (Fig. S2).

**Fig. 2:**
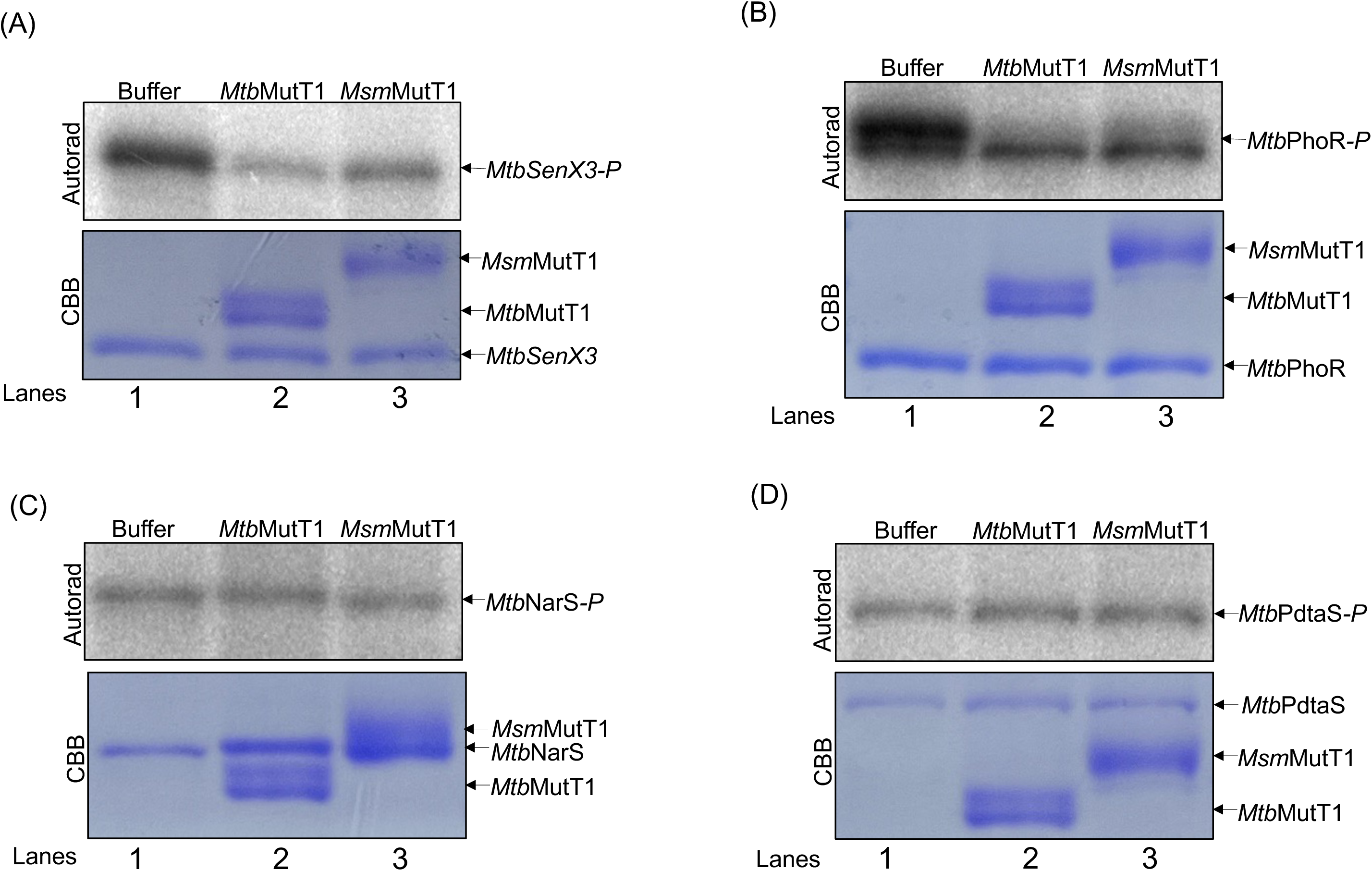
Evaluation of mycobacterial MutT1 phosphatase activity on autophosphorylated SKs. For analysing dephosphorylation, *Mtb*MutT1 or *Msm*MutT1 proteins (1 μg) were mixed with autophosphorylated SKs as follows: **(A)** *Mtb*SenX3-*P* (1 μg); **(B)** *Mtb*PhoR*-P* (1 μg); **(C)** *Mtb*NarS*-P* (0.5 μg); **(D)** *Mtb*PdtaS*-P* (0.5 μg). The reactions were incubated for 1 h at 30^°^C and the gels were autoradiographed. The top panels in **A-D** are autoradiograms (Autorad) and bottom panels are the corresponding Coomassie Brilliant Blue (CBB) stained gels. The images presented are representative of three independent experiments. It may be noted that due to no heating of the reactions in SDS-PAGE sample buffer, MutT1 proteins migrate as doublets.

**Fig. 3:**
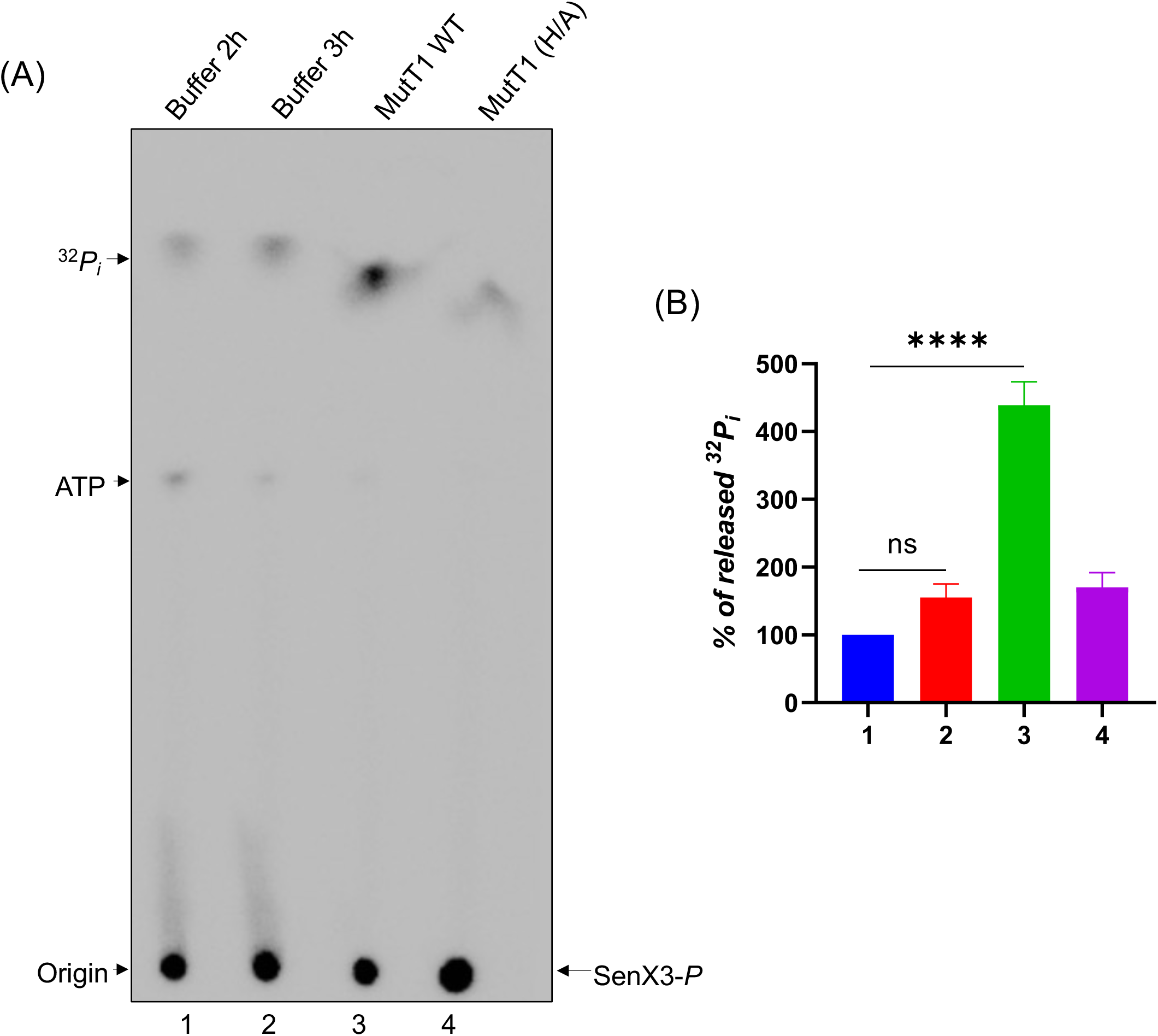
Investigation of *P_i_* release upon MutT1 mediated dephosphorylation of SK-*P*. **(A)** Autophosphorylated SenX3 (1 μg) was incubated with buffer alone or with 2 μg each of the wildtype or phosphatase dead MutT1 for 1 h at 30^°^C. The samples were spotted and separated on PEI Cellulose F plate and autoradiography was done to record the ^32^*P_i_* release. **(B)** Bar graph showing % of released ^32^*P_i_* quantified from panel **(A)**. Bars represent mean ± SD for n = 3. One-way ANOVA was used to calculate *P* values. ∗∗∗∗ <0.0001 indicate significant differences between samples; ns- represent not significant.

### The RHG motif in MutT1 CTD is crucial for its phosphatase activity on SKs

To elucidate which of the MutT1 domains is responsible for the phosphatase activity on SKs, we generated single residue mutations in the specific motifs of *Mtb*MutT1 and *Msm*MutT1. Earlier studies have shown that the E, and H residues are crucial for the catalytic activities mediated by the Nudix box, and RHG histidine phosphatase domains, respectively [37–39]. The mutations, E69A targeting the Nudix hydrolase motif, and H161A targeting the RHG motif were introduced in *Mtb*MutT1 (Fig. 4A). The corresponding mutations for *Msm*MutT1, were E81A and H170A (Fig. 4B). To avoid any endogenous contaminating MutT1 activity, the mutant proteins were purified from *M. smegmatis mc^2^155ΔmutT1*::*hyg* strain deleted for *mutT1* gene (Fig. S3). As shown in Fig. 4C, incubation of *Mtb*PhoR-*P* with *Mtb*MutT1 or *Mtb*MutT1 (E69A) resulted in a pronounced reduction of *Mtb*PhoR-*P* levels (Fig. 4C, autoradiogram, compare lanes 2 and 4, respectively). However, no changes in the level of *Mtb*PhoR-*P* occurred when it was incubated with *Mtb*MutT1 H161A mutant (Fig. 4C, autoradiogram, compare lane 3 with buffer alone lane 1). The CBB stained gel confirms that equal amounts of PhoR protein (Fig. 4C) alongside the quantification of the phosphorylated forms of *Mtb*PhoR (Fig. 4D). Essentially, identical results were obtained when *Mtb*PhoR-P was treated with *Msm*MutT1 or its mutants (Figs. 4E and 4F). Use of another SK, SenX3 also presented with the same observations when *Mtb*SenX3-P was treated with *Mtb*MutT1 or its mutants (Figs. 5A and 5B); or when *Msm*SenX3-P was treated with *Msm*MutT1 or its mutants (Figs. 5C and 5D). We note that MutT1 often migrates as a doublet, especially in these SDS-PAGE where samples are loaded without heating in the sample buffer, necessary for studying SK phosphorylation [35].

**Fig. 4:**
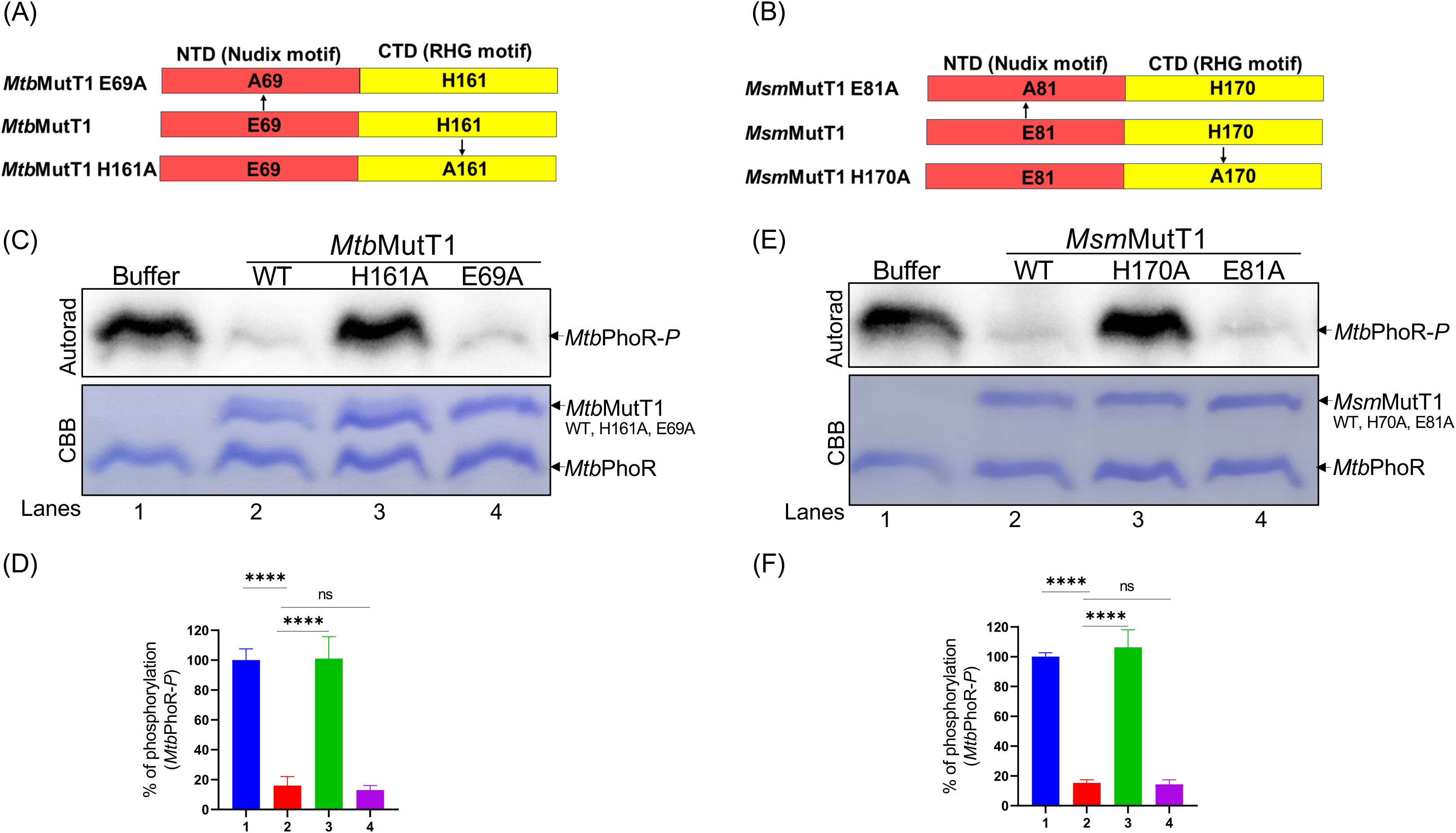
Evaluation of domain specificity in MutT1 mediated SK *Mtb*PhoR-P dephosphorylation. A schematic depicting **(A)** *Mtb*MutT1 **(B)** *Msm*MutT1 with its two domains (NTD and CTD), and the positions where mutations were introduced to inactivate either the Nudix box hydrolase activity in NTD or the RHG histidine phosphatase activity in CTD. **(C)** The dephosphorylation assay was performed by incubating 1 μg of autophosphorylated SK *Mtb*PhoR*-P* with either buffer alone or equal amounts of wildtype and mutant (H161A and E69A) *Mtb*MutT1. (**D**) Densitometric analysis of panel **(B)** showing % phosphorylation levels of *Mtb*PhoR. **(E)** The dephosphorylation assay was performed by incubating 1 μg of autophosphorylated SK *Mtb*PhoR*-P* with either buffer alone or equal amounts of wildtype or mutant (H161A or E69A) *Msm*MutT1. (**F**) Densitometric analysis of panel **(E)** showing % phosphorylation levels of *Mtb*PhoR. The top panels in **(C)** and **(E)** are autoradiograms and bottom panels are the corresponding CBB stained gels. Bar graphs **(D)** and **(F)** represent mean ± SD for n = 3. *P* values were calculated using One-way ANOVA ∗∗∗∗ <0.0001 indicate significant differences between the samples; ns- represent not significant.

**Fig. 5:**
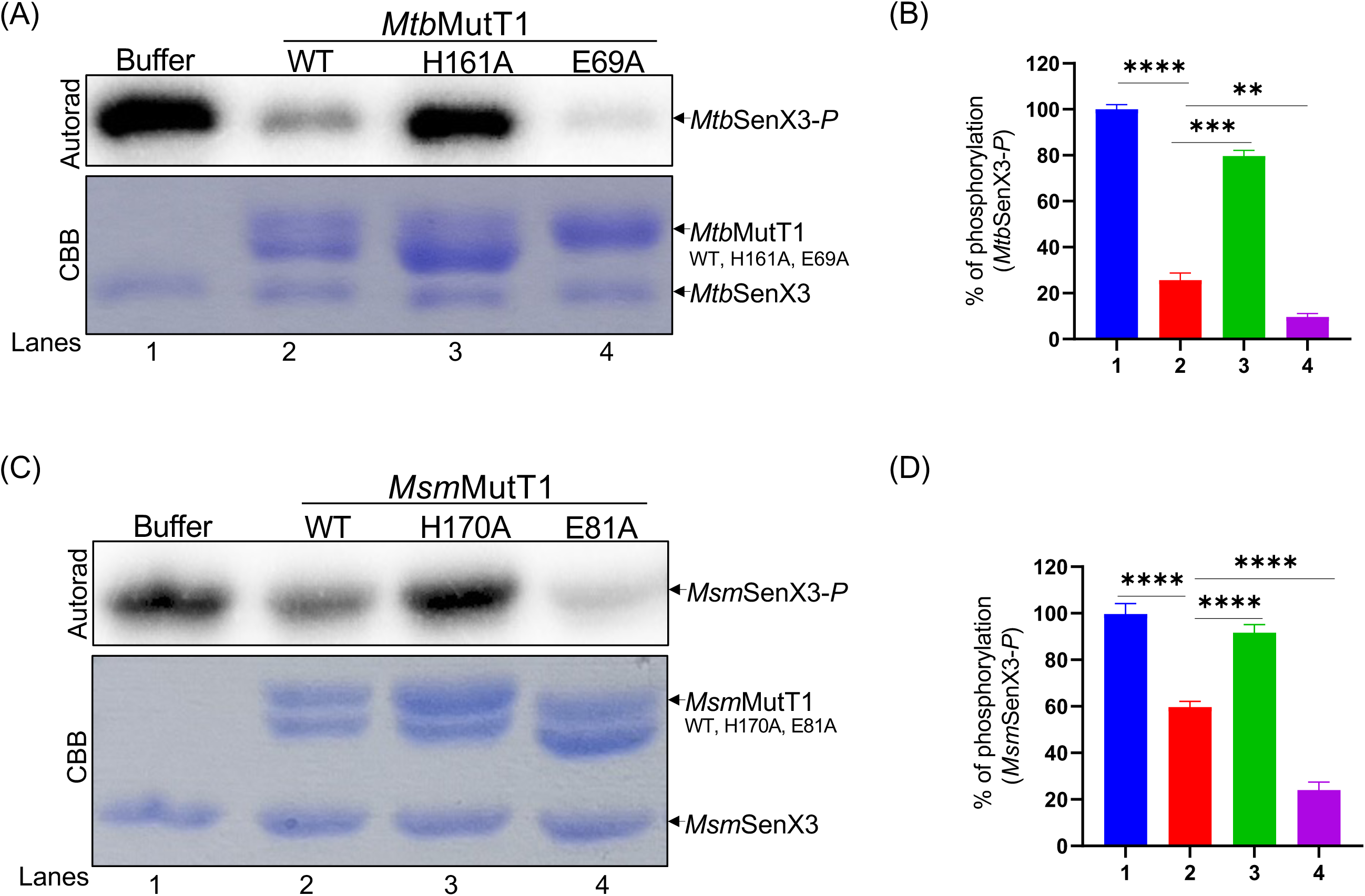
Evaluation of domain specificity in MutT1 mediated SK SenX3-P dephosphorylation. **(A)** The dephosphorylation assay was performed by incubating 1 μg of autophosphorylated SK *Mtb*SenX3*-P* with either buffer alone or equal amounts of wildtype and mutants (H161A and E69A) *Mtb*MutT1. **(B)** Densitometric analysis of panel **(A)** showing % phosphorylation levels of *Mtb*SenX3*-P*. **(C)** 1 μg of autophosphorylated SK *Msm*SenX3*-P* was incubated with either buffer alone, or equal amounts of wildtype or mutants (H161A or E69A) *Msm*MutT1. **(D)** Densitometric analysis of panel **(C)** showing % phosphorylation levels of *Msm*SenX3*-P*. The top panels in **(A)** and **(C)** are autoradiograms and bottom panels are the corresponding CBB stained gels. Bar graphs **(B)** and **(D)** represent mean ± SD for n = 3. *P* values were calculated using One-way ANOVA ∗∗P < 0.01; ∗∗∗P < 0.001; ∗∗∗∗ <0.0001 indicate significant differences between the samples; ns- represent not significant.

Taken together, these findings support the pivotal role of the RHG motif of MutT1 CTD in mediating dephosphorylation of autophosphorylated SKs. It may also be noted that the E69A and E81A mutants in the Nudix boxes of *Mtb*MutT1 and *Msm*MutT1, respectively, appear to have slightly higher phosphatase activity, indicating a possible regulation of the MutT1 CTD phosphatase activity on SenX3 by the MutT1 NTD (Figs. 5A and 5C, compare lanes 4 with 1 in the respective autoradiogram).

### Small molecules like 8-oxo-GTP, GTP and GDP may regulate the activity of MutT1 CTD

*Msm*MutT1 (E81A) exhibited an enhanced activity compared to *Msm*MutT1 (wilt type, WT). The enhancement may result from a longer residency of the substrates (such as 8-oxo-GTP or 8-oxo-dGTP) in the active site pocket of the E81A mutant (with the active site residue mutated) [33]. Thus, we investigated the dephosphorylation status of *Msm*SenX3-*P* in 5 min reactions where clear impact of 8-oxo-GTP could be assessed. No significant changes in the phosphorylation levels of *Msm*SenX3-*P* were detected, when it was incubated alone or with 8-oxo-GTP. There was minimal dephosphorylation with *Msm*MutT1 which was enhanced by the presence of 8-oxo-GTP but inhibited by the presence of 8-oxo-GDP (Fig. S4A, autoradiogram). We then examined the effects of GTP and GDP on *Msm*SenX3-*P* dephosphorylation by MutT1 in 5-, 10-, and 15-min reactions. Higher reduction in the phosphorylation level of *Msm*SenX3*-P* occurred with *Msm*MutT1 in the presence of 8-oxo-GTP, GTP or GDP (compared to *Msm*MutT1 alone) (Fig. S4B autoradiogram). As shown in Figs. 6A and 6B, there is a reduction of ∼30% in the phosphorylation level of *Msm*SenX3-*P* when treated with *Msm*MutT1 (WT); the extent of this reduction increased when 8-oxo-GTP was included along with *Msm*MutT1 (WT). Interestingly, a similar reduction in the phosphorylation levels of *Msm*SenX3-*P* occurred with *Msm*MutT1 (E81A) alone. These findings indicate the significance of E81 in regulating the MutT1 activity towards SenX3. Additionally, it supports the notion of regulation of MutT1 activity by nucleotides such as 8-oxo-GTP.

**Fig. 6:**
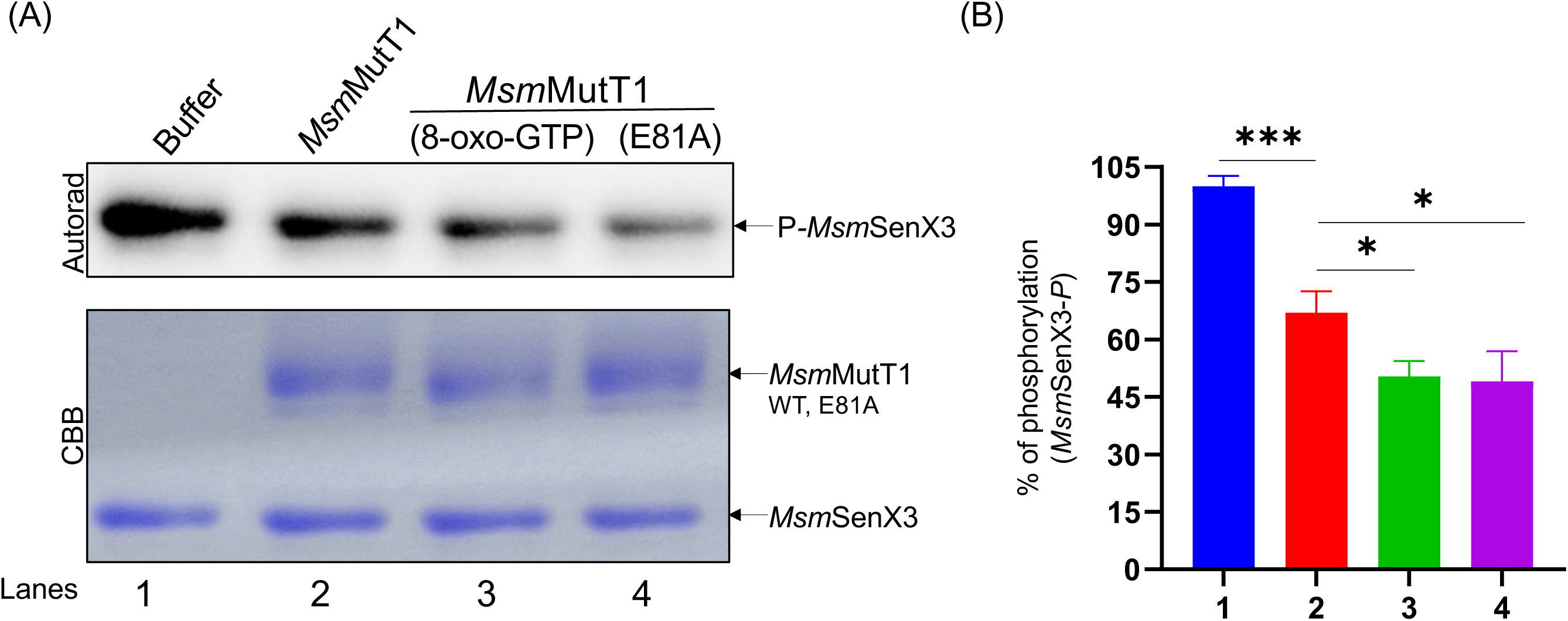
Assessing the impact of 8-oxo-GTP in regulation of MutT1 phosphatase activity. **(A)** The phosphorylated SK *Msm*SenX3 was incubated with either buffer alone, *Msm*MutT1, *Msm*MutT1 + 500 μM 8-oxo-GTP or *Msm*MutT1 (E81A) proteins. The top panel is autoradiogram and bottom panel is the corresponding CBB stained gels. **(B)** Densitometric analysis of panel **(A)** showing % phosphorylation levels of *Msm*SenX3-*P*. Bars represent mean ± SD for n = 3. *P* values were calculated using One-way ANOVA. ∗P < 0.05; ∗∗∗P < 0.001 indicate significant differences between the samples; ns- represent not significant.

### MutT1 impacts phosphotransfer activity from SK to RR

Having demonstrated the activity of MutT1 in dephosphorylating SKs like SenX3-*P*, we sought to investigate its impact on the phosphorylation of its cognate RR, RegX3. We incubated *Msm*SenX3-*P* without or with *Msm*MutT1 or *Msm*MutT1 (H170A), and without or with *Msm*RegX3. As expected, in the reactions with *Msm*SenX3-*P* alone, treatment with *Msm*MutT1 but not with its H170A mutant resulted in ∼40% decrease in the levels of *Msm*SenX3-*P* (Figs. 7A and 7B, compare lane 3 with 1 or 5). Incubation of *Msm*RegX3 with *Msm*SenX3-*P* led to phosphotransfer from *Msm*SenX3-*P* to *Msm*RegX3 (Fig. 7A, autoradiogram, compare lanes 1 and 2). However, in the presence of *Msm*MutT1, the level of *Msm*RegX3-*P* decreased (Figs. 7A and 7C, compare lanes 4 and 2). Interestingly, a slight reduction in transfer of phosphate from SenX3-*P* to RegX3 occurred even in the presence of H170A mutant of *Msm*MutT1 (Figs. 7A and 7C, compare lanes 2 and 6), indicating that even though the H170A mutant is inactive in dephosphorylating SenX3-*P*, it can still impair phosphotransfer from SenX3-*P* to RegX3.

**Fig. 7:**
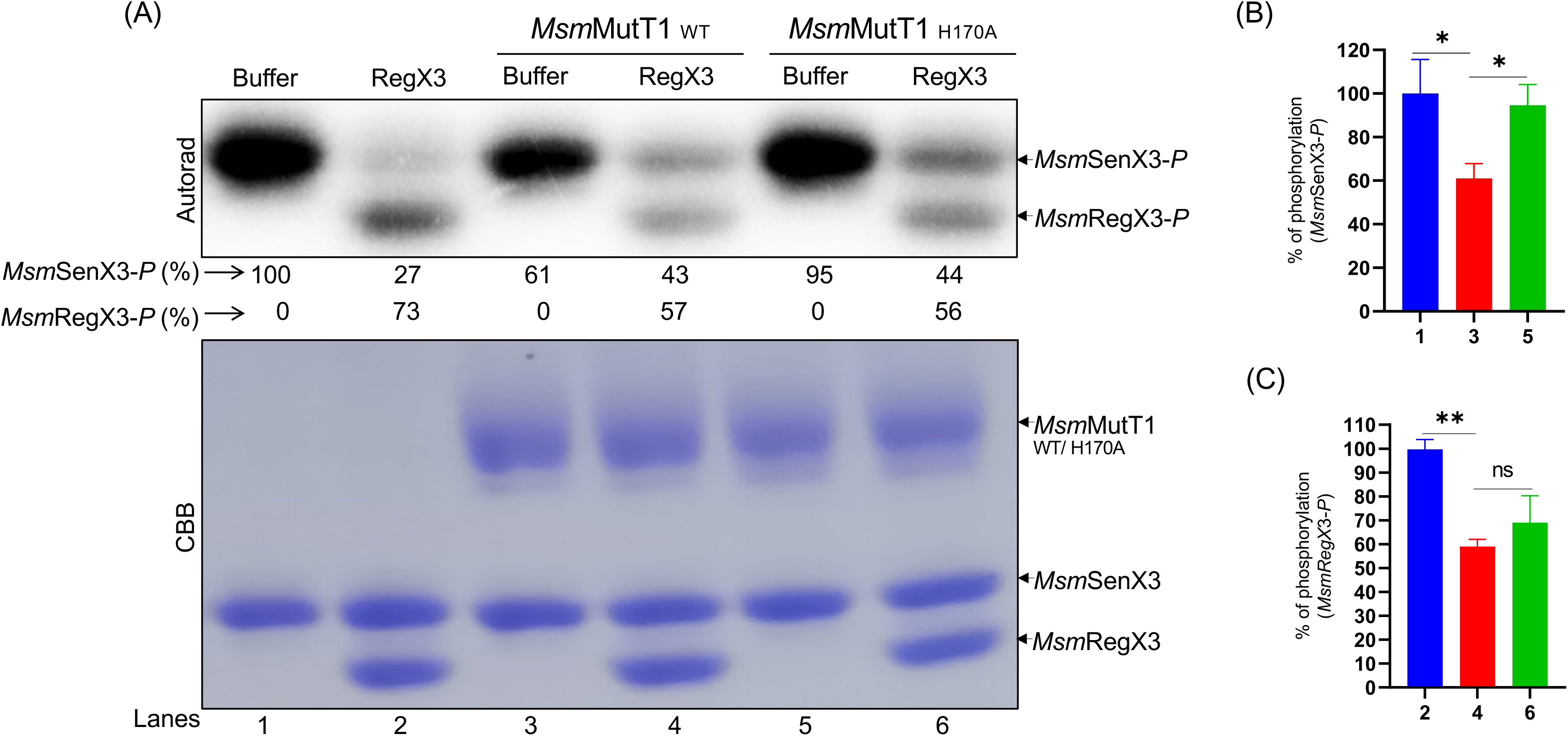
Impact of MutT1 on phosphotransfer from SK-*P* to RR. (**A**) SK SenX3 was autophosphorylated and then incubated with either buffer alone, wildtype or H170A mutant of *Msm*MutT1 for 1h at 30^°^C. For phosphotransfer, the reaction mixture were further incubated with RR RegX3 for 5 min at 30 °C and autoradiography was performed (see methods section). The top panel is autoradiogram and bottom panel is the corresponding CBB stained gels. (**B**) and **(C)** Densitometric analysis from Fig. A showing % phosphorylation levels of SenX3 and RegX3, respectively. Bars represent mean ± SD for n = 3. *P* values were calculated using One-way ANOVA. ∗P < 0.05; ∗∗P < 0.01 indicate significant differences between the samples; ns- represent not significant.

### MutT1 exhibits affinity towards phosphorylated or unphosphorylated SKs

To investigate if the phosphorylation status of the SKs influences its interaction with MutT1, we employed microscale thermophoresis (MST) using phosphorylated and unphosphorylated forms of SK, fluorescently tagged *Mtb*SenX3 (*Mtb*SenX3-mClover). Our analysis revealed ∼1.9-fold higher binding affinity between *Mtb*MutT1 and *Mtb*SenX3-mClover with K_D_ = 223.28 ± 98.17 nM in the phosphorylated state of the SK (Fig. 8A) vs K_D_ = 424.99 ± 130.14 nM for the unphosphorylated state of the SK (Fig. 8B). Interestingly, we obtained similar binding affinities when *Mtb*MutT1 H161A mutant was used with *Mtb*SenX3-mClover, with K_D =_ 195.96 ± 96 nM in the phosphorylated state of SK (Fig. 8C) vs K_D_ = 465 ± 99.62 nM in the unphosphorylated state of the SK (Fig. 8D). As a control, we measured the binding affinity of *Mtb*MutT1 with *Mtb*PdtaS-GFP protein (a non-substrate SK for MutT1), and with the fluorescent protein mClover. No detectable binding was observed for either (Figs. S5A and S5B), confirming the specificity of MutT1 towards SKs. To further validate these findings, we assessed the effect of MutT1 (WT and mutants) on the autophosphorylation of the SK, SenX3. Inhibition in the autophosphorylation of SenX3 occurred when incubated with MutT1 (WT) or E81A or H170A mutants (Fig. S5C).

**Fig. 8:**
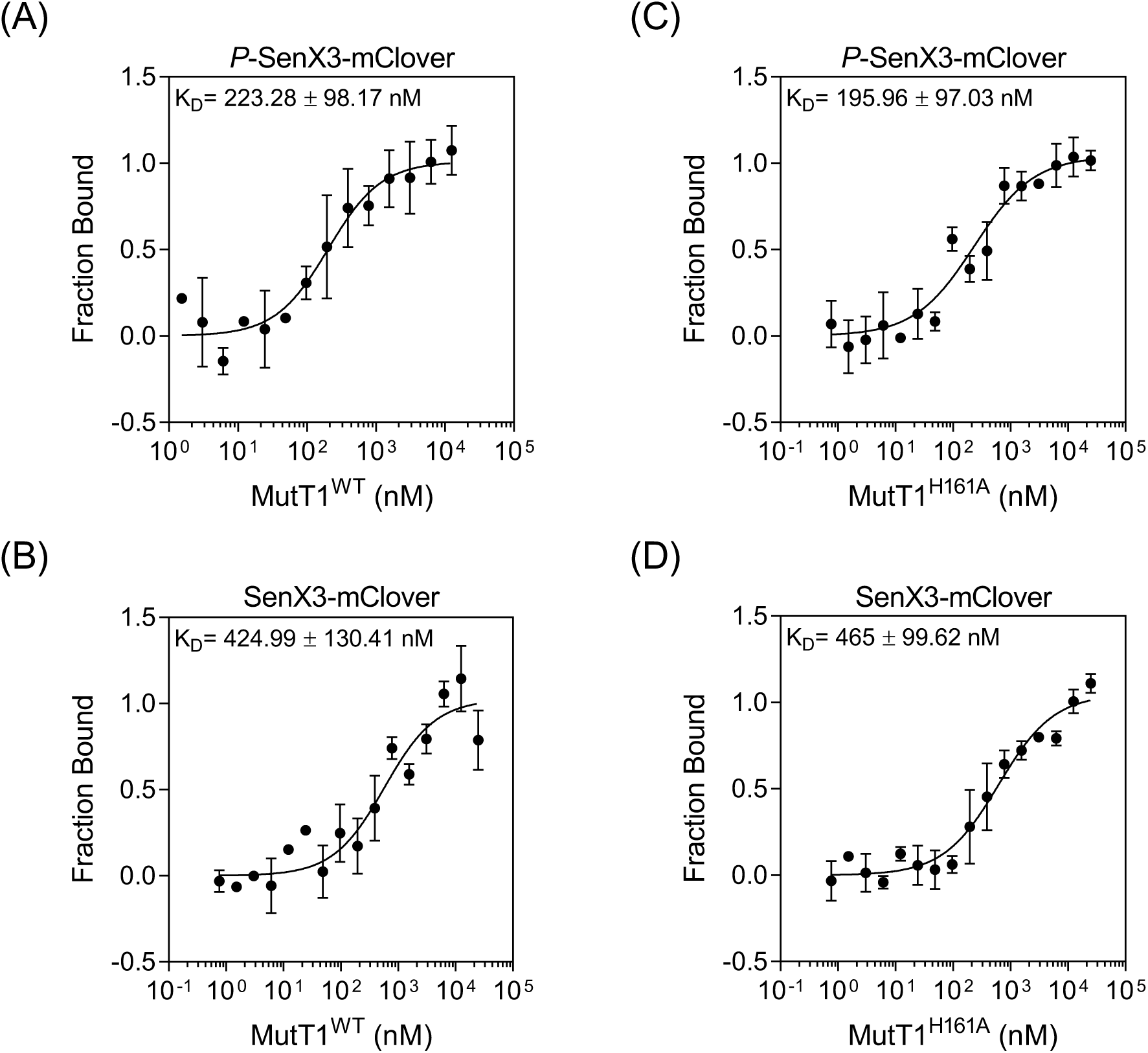
Binding affinities of phosphorylated and unphosphorylated SK SenX3-mClover with MutT1. Normalized fluorescence intensity obtained from MST of 50 nM of the phosphorylated or unphosphorylated SK, SenX3-mClover as a function of the concentration of the titrant MutT1 proteins. **(A)** Phosphorylated SenX3-mClover with MutT1 (concentration range-0.45 nM to 15 μM). (**B**) Unphosphorylated SenX3-mClover with MutT1 (concentration range-0.45 nM to 15 μM). (**C**) Phosphorylated SenX3-mClover with MutT1 H161A mutant (concentration range-0.45 nM to 15 μM). (**D**) Unphosphorylated SenX3-mClover with MutT1 H161A mutant (concentration range-0.45 nM to 15 μM). The resulting K_D_ values are indicated. Curves are best-fits and symbols are mean ± SD (n = 3 independent experiments).

Together, these results demonstrate a reasonably strong affinity of MutT1 with phosphorylated SKs, such as SenX3. The decreases in SK phosphorylation levels by MutT1 or its mutants support its regulatory role in modulating the downstream activities the SKs.

### MutT1 regulates the activity of downstream targets of TCSs

In the context of TCS, the crucial step of regulation involves its impact on the expression of target genes in response to specific environmental conditions. For example, PhoPR is known to be activated, and govern its target gene expression under acidic condition [40]. The gene expression and promoter binding analysis have revealed *lipF* as one of the targets for PhoPR system [41, 42]. Likewise, *prpR* expression was shown to be upregulated by TCS SenX3-RegX3 in MOPS propionate media, particularly when phosphate availability is limited [43]. The regulation is an outcome of interaction between RR, RegX3 and the *prpR* promoter [43]. We selected these targets to investigate the impact of MutT1 on the downstream signaling cascades of TCSs using a *xylE* reporter assay [44, 45]. *M. tuberculosis lipF* and *mutT1* promoters were independently cloned upstream of a promoterless *xylE* reporter gene in pTKmx [45], and introduced into *M. smegmatis* mc^2^155 strain. We observed a significant upregulation of *lipF* promoter activity when cells were grown in acidic media (pH 4.7), compared to cells grown in pH 6.8 media (Fig. S6A). Interestingly, in the same experimental setup there was a significant decrease in the *mutT1* promoter activity under the acidic conditions (Fig. S6B). Given that the role of MutT1 histidine phosphatase is to dephosphorylate PhoR-*P*, the decline in MutT1 promoter activity at pH 4.7 correlates well with the increase in the *lipF* promoter activity, which is positively regulated by activated RR, PhoP. The activity of *prpR* promoter (a target of RR, RegX3) revealed modest increase in the activity when cells were grown in MOPS propionate low phosphate media compared to high phosphate media (Fig. S6C). In the low phosphate medium, there was no significant change in MutT1 promoter activity (Fig. S6D).

Our investigation on the impact of *mutT1* knockout or overexpression on *lipF* and *prpR* promoters revealed intriguing insights. As expected, we observed no discernible differences in *lipF* promoter activity between mc^2^155Δ*mutT1* and mc^2^155 (bars 1 and 2, respectively) at pH 4.7 (Fig. 9A). However, a significant reduction in *lipF* promoter activity occurred when *Msm*MutT1, *Mtb*MutT1, *Msm*MutT1 (E81A), or *Msm*MutT1 (H170A) were overexpressed in the mc^2^155 strain (Fig. 9A, bars 3 to 6). Although, there was a decrease in the *lipF* promoter activity upon overexpression of *Msm*MutT1 (H170A) mutant lacking the PhoR-*P* dephosphorylation activity, the decrease was less compared to the *Msm*MutT1 (Fig. 9A, compare bar 6 to 4). While the observation is consistent with the expected trend, any decreases in *lipF* promoter activity in response to the *Msm*MutT1(H170A) mutant compared with *Msm*MutT1 (WT) suggest presence of additional regulatory mechanisms (*see Discussion*).

**Fig. 9:**
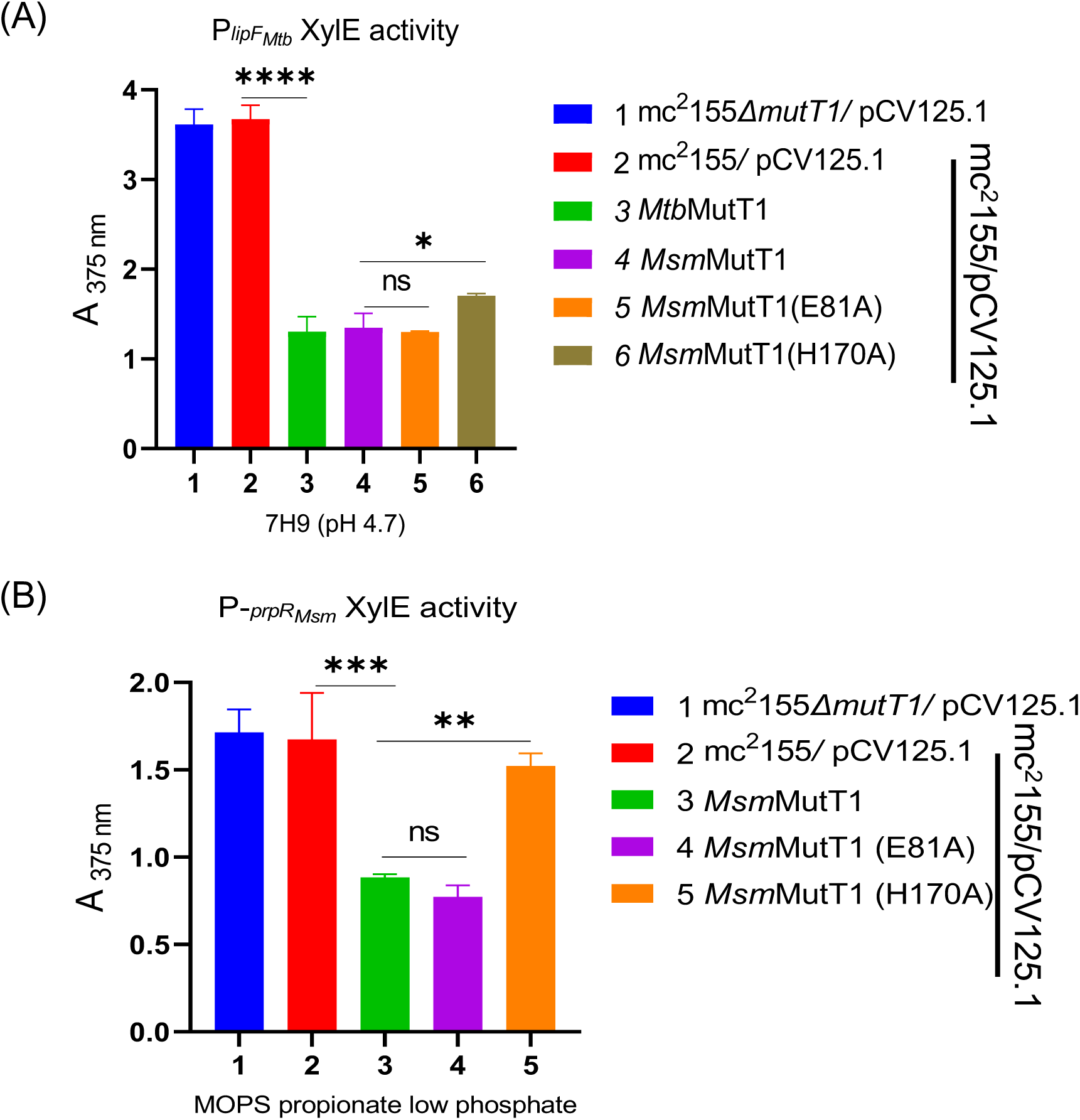
Investigation of wildtype or mutant MutT1 overexpression and its impact on the TCSs target gene expression. The activities of the selected TCSs target gene promoters were analysed using *xylE* reporter. **(A)** *lipF* promoter (a target of TCS PhoPR in *Mtb*) was introduced upstream of *xylE* ORF. The resulting reporter construct were electroporated in the following *M. smegmatis* strains, 1- *Msm* mc^2^155 (Δ*mutT1*)/ pCV125.1; 2- *Msm* mc^2^155 (WT)/ pCV125.1; 3- *Msm* mc^2^155 (WT)/ pCV125.1:*Mtb*MutT1; 4- *Msm* mc^2^155 (WT))/ pCV125.1:*Msm*MutT1; 5- *Msm* mc^2^155 (WT)/ pCV125.1:*Msm*MutT1 E81A; 6- *Msm* mc^2^155 (WT)/pCV125.1:*Msm*MutT1 H170A. Cultures were grown in 7H9 media at pH 4.7 and used to make cell-free extracts for XylE assays. **(B)** *prpR* promoter (a target of TCS SenX3-RegX3 in *Msm*) was introduced upstream of *xylE* ORF. The resulting reporter construct was electroporated in the following *M. smegmatis* strains, 1- *Msm* mc^2^155 (Δ*mutT1*)/ pCV125.1; 2- *Msm* mc^2^155 (WT)/ pCV125.1; 3- *Msm* mc^2^155 (WT))/ pCV125.1:*Msm*MutT1; 4- *Msm* mc^2^155 (WT)/pCV125.1:*Msm*MutT1 E81A; 5- *Msm* mc^2^155 (WT)/ pCV125.1:*Msm*MutT1 H170A. Cultures were grown in MOPS propionate media with low phosphate concentration (100 μM) and used to make cell-free extracts. Bars represent mean ± SD for n = 3. One-way ANOVA method was used to calculate *p* values. ∗ *p* < 0.05; ∗∗ *p* < 0.01; ∗∗∗ *p* < 0.001; ∗∗∗∗ *p* < 0.0001 indicate significant differences between samples; ns- represent not significant.

Next, we assessed the impact of *mutT1* knockout or overexpression on the promoter activity of *prpR* in MOPS propionate low phosphate medium. We observed no differences in the *prpR* promoter activity in mc^2^155Δ*mutT1* and mc^2^155 (Fig. 9B, bars 1 and 2, respectively). Upon overexpression of *Msm*MutT1 and *Msm*MutT1(E81A), there was a clear decrease in the *prpR* promoter activity (Fig. 9B, bars 3 and 4, respectively). However, a very small but significant decrease in the *prpR* promoter activity was also observed upon overexpression of the phosphatase dead *Msm*MutT1 (H170A) mutant (Fig. 9B, bar 5). In yet another case study, we checked for the impact of MutT1 expression on the promoter activity of *kdpA*, a target of TCS KdpED [46, 47]. We showed that SK KdpD-*P* is not a substrate for MutT1 (Fig. S1K). Nonetheless, we observed a decrease in the *kdpA* promoter activity upon *Msm*MutT1 expression (Fig. S7A). To check the specificity of the effect, we used a reporter driven by *hsp60* promoter, which remained largely unchanged upon MutT1 expression (Fig. S7B). In fact, the activity of *hsp60* promoter remained unchanged even at a lower pH of the medium (Fig. S7C).

Clearly, these specific examples of the downstream target genes show the physiological impact of MutT1 on TCSs mediated gene regulation. The findings reveal the role of MutT1 in regulating TCSs and their target genes, particularly as exemplified by its impact on *lipF* and *prpR* promoters. However, it should also be said that the *in vivo* impact of dysregulating TCSs is complex and the outcomes of perturbing an isolated element/player in the pathway are not always straightforward. For example, in the case of KdpA, the impact of MutT1 might have been mediated through dephosphorylation of SK MtrB-*P*, which is known to cross-regulate RR KdpE [30]. These observations, while revealing a regulatory role of MutT1, underscore the complex nature of interplay between MutT1 and its outcome on TCSs.

## Discussion

Drawing from the structural similarities between the mycobacterial MutT1 CTD and *E. coli* SixA, and the presence of RHG motif in both the proteins (Fig. 1B) [33], we hypothesized that MutT1 may share the regulatory roles of SixA protein. This conjecture was founded on the proposed role of SixA in dephosphorylation of a SK ArcB [36]. Our study provides important evidence of the regulation of a subset of mycobacterial TCSs by MutT1. We have established that MutT1 CTD mediates dephosphorylation of selected SKs through its RHG motif. Moreover, we demonstrated the impact of MutT1 on the downstream activity of TCS.

MutT1 exhibits an unprecedented phosphatase activity towards a subset of SKs such as PhoR, SenX3, MtrB, MprB, DosS and PrrB (Figs. 2A and 2B; Figs. S1A-E); but not the ones such as NarS, PdtaS, KdpD, DosT and TcrY (Figs. 2C and 2D; Figs. S1F-K). These findings highlight the specificity of the phosphatase activity of MutT1 on SKs. Further, the mutational analyses revealed that mutation in Nudix hydrolase motif does not abolish the MutT1 phosphatase activity towards the selected SKs. In contrast, mutation in the RHG motif resulted in a loss of MutT1 phosphatase activity on the SKs (Figs. 4 and 5). These data reveal the crucial role of the RHG motif in mediating MutT1 phosphatase activity on SKs, corroborating the structural comparisons between *Eco*SixA and the CTD of *Msm*MutT1 [33]. Interestingly, the Nudix hydrolase motif mutations (E69A in *Mtb*MutT1 and E81A in *Msm*MutT1) appeared to enhance the MutT1 phosphatase activity on SK, SenX3-*P* (Fig. 5). This indicates a potential regulatory role of NTD in modulating the histidine phosphatase activity of the CTD, particularly concerning the SenX3-*P*. MutT1 is known for its activity towards small molecules like 8-oxo-dGTP or 8-oxo-GTP, which come in close contact with E81 residue in the Nudix hydrolase motif of *Msm*MutT1 [33]. We hypothesized that these molecules may regulate the activity of MutT1 CTD on SK dephosphorylation. Our data show that small molecules like 8-oxo-GTP, GTP and GDP enhance the activity of MutT1 on SenX3-P (Fig. 6; Fig. S4B).

Following autophosphorylation of SKs, the transfer of the phosphate group from its His residue to the Asp residue of its cognate RR is responsible to regulate the target genes expression. Consistent with the MutT1 mediated dephosphorylation of the SKs, we observed a conspicuous reduction in the phosphorylation level of RegX3 upon incubation with SenX3-*P* with MutT1 or MutT1 (H170A) (Fig. 7A to 7C). These data suggest that, although MutT1 (H170A) is inactive for its SixA like activity, it may still interact with the SKs, sequestering it and inhibiting its phosphotransfer activity. These observations were further validated when the binding affinity of MutT1 WT and mutant with SenX3 was evaluated (Fig. 8).

The impact of MutT1 on the expression of the selected target genes revealed a substantial reduction in the activity of *lipF* promoter by MutT1 and MutT1 (E81A) overexpression. However, to our surprise, while assessing the impact of MutT1 overexpression, the expression from the *lipF* promoter was still higher upon MutT1 (H170A) expression, it was considerably lower than the controls suggesting a complex mode of regulation by RR, PhoP. In fact, a recent study shows that PhoP mediated regulation is also impacted by the cAMP levels [48]. While a detailed analysis of this observation is necessary, to a first approximation, we believe that this may well be because of the sequestration of SK, PhoR by MutT1 or its mutants when expressed under acidic condition (pH 4.7) (Fig. 9A). Nonetheless, our observations with the *prpR* promoter reveal a significant reduction in its activity in the presence of *Msm*MutT1 or its NTD mutant (E81A) but not with its CTD mutant (H170A) (Fig. 9B).

Of particular interest is the regulation of *kdpA* promoter by MutT1. We note that the SK KdpD is not a substrate for MutT1. Still there is a remarkable decrease in *kdpA* promoter activity upon MutT1 expression (Fig. S7A). Use of *hsp60* promoter as a control yielded no discernible effect of MutT1 expression on its activity (Fig. S7B), suggesting that the effect of MutT1 on *kdpA* promoter is a specific one. This finding is not only consistent with the reports of crosstalk between TCSs [4], but also adds another evidence supporting the role of crosstalk between the SKs and ‘noncognate’ RRs, and the complex nature of TCS mediated regulation. It has been reported that the SK MtrB, which is a substrate for dephosphorylation by MutT1, exhibits phosphotransfer activity towards non-cognate RRs such as PhoP, NarL, KdpE, TcrX and TcrA [30], [49].

In most TCS, positive autoregulation enhances its sensitivity to the input signals. Positive autoregulation is preferable when the signal is strong. However, a positive autoregulation in response to a weak/transient signal may trigger a disproportionate response which may compromise the bacterial fitness, especially in stressful conditions like oxidative stress [8]. To our knowledge, phosphatases acting on the SKs to dampen the TCSs downstream activities, have not been reported in mycobacteria. This report of MutT1 acting as a phosphatase for SKs fills this lacuna to offer a finer regulation of the mycobacterial TCSs.

In light of our study, we propose a general regulatory model elucidating the interplay between MutT1 and the TCSs in mycobacteria. In the absence of MutT1, a SK undergoes autophosphorylation and subsequently regulates the activity of its partner RR by transferring phosphoryl group onto its conserved Asp residue (Fig. 10A). The activated RRs then binds to the promoter region of its target genes, thereby modulating its activity in response to stress. The presence of MutT1 dephosphorylates the phosphorylated SKs, inhibiting its ability to fully activate its partner RR, thereby preventing unnecessary signal responses (Fig. 10B).

**Fig. 10:**
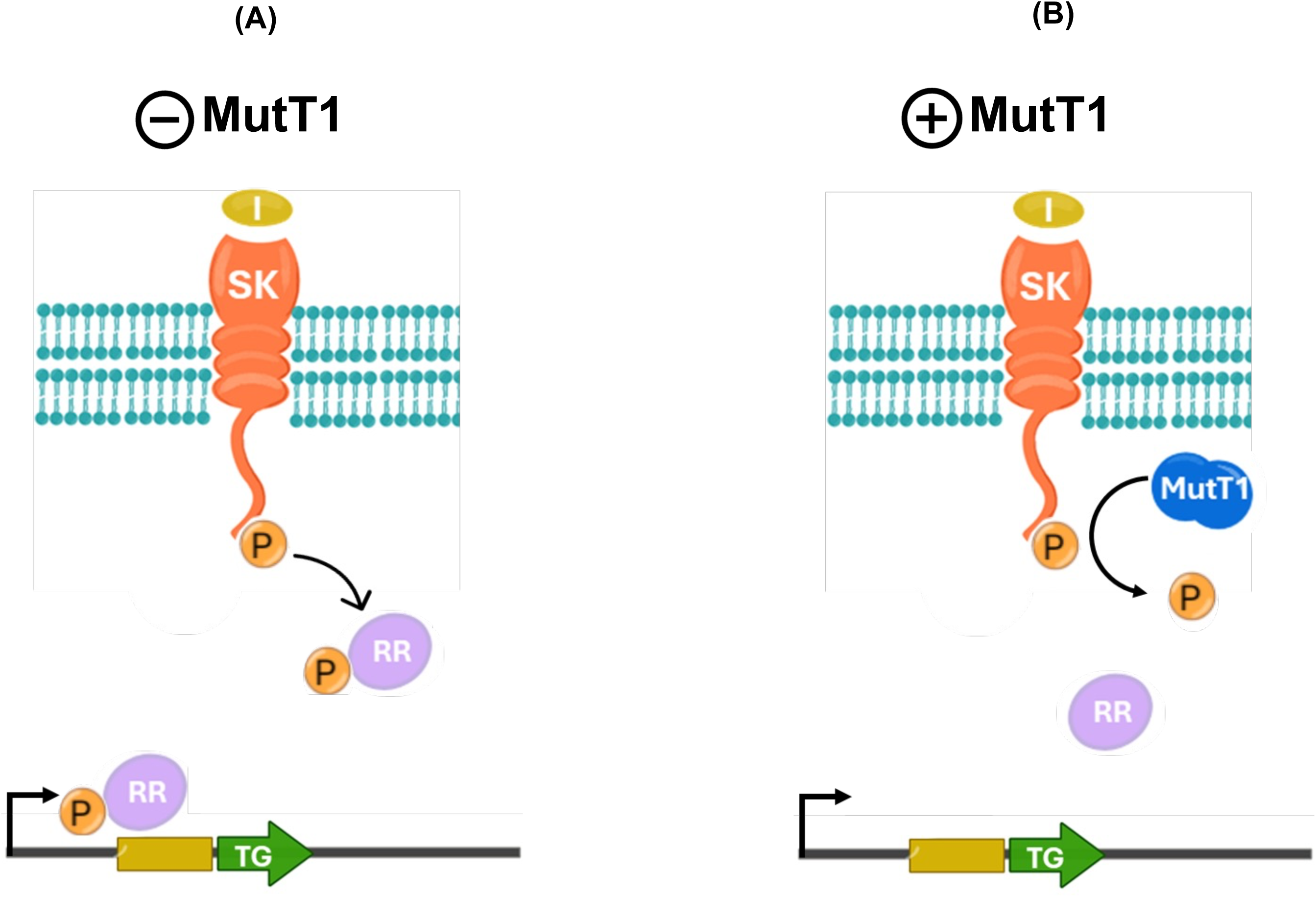
A model showing MutT1 mediated regulation of TCS signaling in bacteria. The schematics are showing a model for MutT1 mediated regulation TCSs phospho-signaling in bacteria. **(A)** Under the environmental condition wherein MutT1 protein is not expressed or functionally inactive, upon signal perception, SKs can autophosphorylate and transfer the phosphoryl group to its partner RRs which can bring downstream changes by binding to its target gene promoters. (B) Under the environmental conditions wherein the MutT1 protein is expressed or functionally active, it can dephosphorylate the activated SKs, thereby reducing the downstream signaling cascade which is operated through its partner RRs.

Our observations shed light on the intricate molecular interplay governing bacterial TCSs and emphasize the regulatory role of MutT1 in maintaining SKs activities. Further elucidation of the mechanistic details underlying this phenomenon could offer valuable insights into the modulation of bacterial cellular responses and adaptive behavior. Moreover, our study presents an exciting avenue for potential drug discovery efforts aimed at mimicking the inhibitory effect of MutT1, thereby potentially restraining the pathogen growth.

## Materials and Methods

### Bacterial strains, plasmids, and DNA oligomers

Strains and plasmids used in this study are described in Table 1. DNA oligomers are listed in Table S1. Enzymes used for cloning were ordered from New England Biolabs and Thermo Fisher Scientific.

**Table 1:** Description of strains and plasmids used in the study.

| Strain/Plasmid | Genotype | Reference/Source |
| --- | --- | --- |
| pJAM His-tag | Shuttle vector with ColE1 (*E. coli*) and pAL5000 (mycobacteria) origins of replication, *kan<sup>R</sup>*, and mycobacterial acetamidase promoter | [53] |
| pJAM_ *Msm*MutT1 | *Msm*MutT1 was cloned into pJAM_His between BamHI and XbaI sites. | This study |
| pJAM *Msm*MutT1 E81A | pJAM with *Msm* mutT1 E81A mutant | This study |
| pJAM *Msm*MutT1 H170A | pJAM with *Msm* mutT1 H170A mutant | This study |
| pJAM *Mtb*MutT1 | *Mtb* mutT1 was cloned into pJAM His in XbaI site. | This study |
| pJAM *Mtb*mutT1 E69A | pJAM with *Mtb* mutT1 E69A mutant | This study |
| pJAM *Mtb*mutT1 H161A | pJAM with *Mtb* mutT1 H161A mutant | This study |
| pPROEX: *Msm*SenX3 | *Msm* senX3 gene was cloned into pPROEX HTb between NcoI and HindIII sites. | This study |
| pPROEX: *Msm*regX3 | *Msm* regX3 gene was cloned into pPROEX HTb between NcoI and HindIII sites. | This study |
| pCV125 | An integrating vector (*kan<sup>R</sup>*) and a promoterless *lacZ* gene | [54] |
| pCV125.1 | The kanamycin cassette in the pCV125 vector was replaced with the chloramphenicol cassette between (NheI and SpeI) | This study |
| pCV125.1:*Msm*MutT1(Cm<sup>R</sup>) | Mycobacterial hsp60 promoter + *Msm* mutT1 was digested from pMV261 *Msm*MutT1 with ( XbaI & NdeI) and subcloned into pCV125.1 | This study |
| pCV125.1:*Msm*mutT1 E81A(Cm<sup>R</sup>) | pCV125.1 with *Msm* mutT1 E81A mutant | This study |
| pCV125.1:*Msm*mutT1 H170A(Cm<sup>R</sup>) | pCV125.1 with *Msm* mutT1 H170A mutant | This study |
| pCV125.1:*Mtb*mutT1 (Cm<sup>R</sup>) | Mycobacterial hsp60 promoter + *Mtb* mutT1 was digested from pMV261 *Msm* mutT1 with ( XbaI & NdeI) and subcloned into pCV125.1 | This study |
| pTKmx | *E. coli*-mycobacteria shuttle vector containing kanamycin resistance marker and promoterless *xylE* reporter gene | [45] |
| pTKmx:P<sub>*Mtb* lipF</sub> | *Mtb* lipF promoter cloned upstream of the *xyl* E cassette between the EcoRI and BamHI sites | This study |
| pTKmx:P<sub>*Msm*prpR</sub> | *Msm* prpR promoter cloned upstream of the *xyl* E cassette between the EcoRI and BamHI sites | This study |
| pTKmx:P<sub>*Mtb*mutT1</sub> | *Mtb* mutT1 promoter cloned upstream of the *xyl* E cassette between the EcoRI and BamHI sites | This study |
| pTKmx:P<sub>BGGhsp60</sub> | Mycobacterial hsp60 promoter cloned upstream of the *xyl* E cassette between the EcoRI and BamHI sites | This study |
| pTKmx:P<sub>*Msm*kdpA</sub> | *Msm*kdpA promoter cloned upstream of the *xyl* E cassette between the EcoRI and BamHI sites | This study |

### Media, and growth conditions

*E. coli* strains were grown in Luria-Bertani (LB) medium. For growth on the solid surface, 1.6% agar was added to liquid medium. For culturing *M. smegmatis* strains, LB containing 0.2% tween 80 (v/v) (LBT) media was used. For growth on solid media, LB containing 0.04% tween 80 (v/v) was supplemented with 1.5% agar (w/v). MOPS propionate media was prepared by mixing (25 mM MOPS, pH 7.2, 25 mM KCl, 10 mM NaCl, 10 mM (NH_4_)_2_SO_4_, 10 mM FeCl_3_, 2 mM MgSO_4_, 100 mM CaCl_2_, 10 mM C_3_H_5_NaO_2_, 0.02% Tween-80, 100 μM K_2_HPO_4_ for low phosphate and 10 mM K_2_HPO_4_ for high phosphate). The acidic media (pH 4.7) was prepared by adjusting the pH of 7H9 media with HCl. Media were supplemented with 50 mg/ml ampicillin, 50 mg/ml kanamycin, 50 mg/ml hygromycin and 25 mg/ml chloramphenicol as required. Medium components were purchased from BD Difco (Franklin Lakes, NJ).

### Structural comparisons

For analysing the MutT1 protein and structure comparison of mycobacterial MutT1 CTD with *E. coli* SixA protein, the protein structure files were retrieved from Protein Data Bank website. The structures were aligned using Edu PyMol software [50].

### Generation of *M. smegmatis* mc^2^155 Δ*mutT1* strain

The DNA substrate and *M. smegmatis* mc^2^155 Δ*mutT1* strain were prepared as described [51], [52]. *M. smegmatis* mc^2^155 was electroporated with pNIT vector and plated on LBT-Kan (LBT containing 50 mg/ml Kan) and incubated at 37 °C. A single colony was then inoculated in10 ml LBT-Kan and grown under shaking at 37 °C to obtain saturated culture. A 0.5% inoculum of the saturation culture was used to inoculate 10 ml LBT-Kan and grown till the culture OD_600_ of ∼0.4-0.6 was obtained. The culture was induced with 5 mM isovaleronitrile (IVN) and grown further for 3 h. The cells from the culture were pelleted and prepared for electroporation with 100 ng linear DNA (allele exchange substrate). After electroporation, 1 ml LBT was added and cells were kept for recovery for 4 h at 37 °C, and then plated on LBT containing Hyg (150 mg/ml) and Kan (50 mg/ml) and incubated at 37 °C. For confirmation of the knockout, the genomic DNA was isolated and used for PCR using Msm mutT1 primers.

### Cloning of *Msm*MutT1

The open reading frame (ORF) of *Msm*MutT1 (MSMEG_2390) was amplified from the *M. smegmatis* mc^2^155 genomic DNA using the *Msm* mutT1 BamH1 Fp and *Msm* mutT1 Xba1 Rp. The reaction mixture containing 100 ng template DNA, 10 pmols each of the forward and reverse primers, 250 mM dNTPs, Pfu DNA polymerase buffer, and Pfu DNA polymerase. PCR was incubated at 94 °C for 5 min followed by 30 cycles of incubations at 94 °C for 1 min, 63 °C for 30 s, and 72 °C for 2 min, and then 72 °C for 5 min. The PCR product was purified, digested with BamHI and XbaI, cloned into similarly digested pJAM_His plasmid, and confirmed by restriction endonuclease (RE) digestions and DNA sequencing.

### Cloning of *Mtb*MutT1

The ORF of *Mtb*MutT1 (Rv2985) was amplified from *M. tuberculosis* H37Rv genomic DNA using the Mtb mutT1 Xba1 Fp and *Mtb* mutT1 Xba1 Rp. The reaction containing 100 ng template DNA, 10 pmols each of the forward and reverse primers, 250 mM dNTPs, Pfu DNA polymerase buffer, and Pfu DNA polymerase. PCR was incubated at 94 °C for 5 min followed by 30 cycles of incubations at 94 °C for 1 min, 60 °C for 30 s, and 72 °C for 2 min, and then 72 °C for 5 min. The PCR product was purified and digested with XbaI, cloned into similarly digested pJAM_His, and confirmed by RE digestions and DNA sequencing.

### Cloning of *Msm*SenX3

The cytosolic catalytic region of *Msm*SenX3 (MSMEG_0936) was amplified from *M. smegmatis* mc^2^155 genomic DNA using the *Msm* senX3-Ncoi-Fp and *Msm* senX3-Hindiii-Rp. The reaction mixture of 100 ng template DNA, 10 pmols each of the forward and reverse primers, 250 uM dNTPs, 1x Pfu DNA polymerase buffer, and Pfu DNA polymerase. The PCR was incubated at 94 °C for 5 min followed by 30 cycles of incubations at 94 °C for 1 min, 58 °C for 30 s, and 72 °C for 2 min, and then incubated at 72 °C for 5 min. The PCR product was purified, digested with NcoI and HindIII, cloned into similarly digested pProEx-Htb plasmid, and confirmed by RE digestions and DNA sequencing.

### Cloning of *Msm*RegX3

The open reading frame of *Msm* regX3 (MSMEG_0937) was amplified from *M. smegmatis* mc^2^-155 genomic DNA using Msm regX3-Ncoi-Fp and Msm regX3-Hindiii-Rp. The reaction mixture of 100 ng template DNA, 10 pmols each of the forward and reverse primers, 250 μM dNTPs, Pfu polymerase buffer, and Pfu DN polymerase. The PCR reaction was incubated at 94 °C for 5 min followed by 30 cycles of incubations at 94 °C for 1 min, 58 °C for 30 s, and 72 °C for 2 min, and again 72 °C for 5 min. The PCR product was purified, digested with NcoI and HindIII, cloned into similarly digested pProEx-Htb plasmid, and confirmed by restriction enzyme digestions and DNA sequencing.

### Cloning of promoters for XylE assays

The promoters were amplified from *M. smegmatis* mc^2^155 or *M. tuberculosis* H37Rv genomic DNAs using the primers listed in Table S1. The reaction contained 100 ng template DNA, forward and reverse primers (10 pmols each), 250 mM dNTPs, Pfu DNA polymerase buffer, and Pfu DNA polymerase. The PCR was incubated at 94 °C for 5 min followed by 30 cycles of incubations at 94 °C for 1 min, 58 °C for 30 s, and 72 °C for 1 min, and then 72 °C for 5 min. The PCR products was purified, digested with EcoRI and BamHI and cloned into similarly digested pTKmx-XylE plasmid, and confirmed by RE digestions and DNA sequencing.

### Generation of MutT1 mutants

*Msm*MutT1 E81A or H170A, *Mtb*MutT1 E69A or H161A were generated by site-directed mutagenesis (SDM), using pJAM-*Msm*MutT1 and pJAM-*Mtb*MutT1 templates, respectively. The primers, *Msm* mutT1 E81A Fp, *Msm* mutT1 E81A Rp, *Msm* mutT1 H170A Fp, *Msm* mutT1 H170A Rp, *Mtb* mutT1 E69A Fp, *Mtb* mutT1 E69A Rp, *Mtb* mutT1 H161A Fp, *Mtb* mutT1 H161A Rp, were used for SDM. The reaction with 100 ng template DNA, 10 pmols each of the forward and reverse primers, 250 μM dNTPs, 1x Q5 DNA polymerase buffer, and Q5 DNA polymerase (NEB #M0491) was heated at 98 °C for 1 min followed by 20 cycles of incubations at 98 °C for 20 s, 68 °C for 30 s, and 72 °C for 6 min and final extension at 72 °C for 10 min. The PCR product was digested with DpnI (Thermo Scientific™ # ER1702) and used to transform *E. coli* TG1 cells. Plasmids from the transformants were confirmed for SDM by DNA sequencing.

### Purification of MutT1 proteins

MutT1 proteins were overexpressed and purified from *M. smegmatis* mc^2^155Δ*mutT1*. Briefly, the MutT1 plasmids were electroporated into the mc^2^155Δ*mutT1* strain. A single colony was inoculated in LBT-Kan and grown under shaking at 37 °C to make saturated culture. Inoculum (1%) of the saturated culture was sub-cultured into 3000 ml LBT-Kan. At OD_600_ of ∼0.6, expression of the proteins was induced with 1% acetamide and the cultures were grown further at 30 °C for 9 h. The cells were pelleted by centrifugation, resuspended in buffer A (20 mM Tris-HCl, pH 8.0, 1 M NaCl, 10% (v/v) glycerol, 20 mM imidazole and 2 mM β-mercaptoethanol), lysed by sonication and subjected to ultracentrifugation at 26K rpm at 4 °C for 2 h (using optima XPNs Ultra Centrifuge, Beckman Coulter). The clarified lysate was loaded onto pre-equilibrated Ni-NTA column. The column was washed with washing buffer (1 M NaCl, 20 mM Tris-HCl pH 8, 10% (v/v) glycerol, 2 mM β-mercaptoethanol and 40 mM imidazole). The proteins were eluted with a gradient of 40 to 1000 mM imidazole in wash buffer lacking imidazole. The fractions containing the desired protein were pooled, concentrated, and loaded onto Superdex-75 column, and eluted with gel filtration buffer (1 M NaCl, 20 mM Tris-HCl pH 8, 10% (v/v) glycerol and 2 mM β-mercaptoethanol). The fractions containing the purified proteins were pooled and dialyzed against dialysis buffer (750 mM NaCl, 20 mM Tris-HCl pH 8, 10% (v/v) glycerol, and 2 mM β-mercaptoethanol), followed by dialysis in storage buffer (500 mM NaCl, 20 mM Tris-HCl pH 8, 50% glycerol, and 2 mM β-mercaptoethanol) and stored at −20 °C.

### Purification of cytosolic kinase domain of SKs and full-length RR proteins

The proteins were over-expressed and purified from *E. coli* strains carrying expression plasmids were grown at 37 °C in 1000 ml 2xYT until OD_600_ of ∼0.4−0.6. IPTG (500 mM) was added and the culture was grown further for 15–20 h at 11 °C for protein expression. Cells were harvested by centrifugation, and purified as described previously [49].

### SK dephosphorylation assay

Briefly, 1 mg of SK was incubated in kinase buffer (50 mM Tris-HCl, pH 8.0, 50 mM KCl, 10 mM MgCl_2_) containing 1 μCi of γ-^32^P-labelled ATP at 30 °C for 1 h, and then incubated with 1 to 2 mg MutT1 proteins for for indicated timepoints at 30 °C. The reaction was terminated by adding 1× SDS-PAGE sample buffer (heating step was not included). The samples were resolved on 12% SDS-PAGE, and exposed to phosphor screen (Fujifilm Bas cassette2, Japan) for 4 h and imaged using Typhoon 9210 phosphor imager (GE Healthcare, USA).

### Regulation of SenX3 with small molecules

Briefly, 1 μg SenX3 was incubated in kinase buffer (50 mM Tris-HCl, pH 8.0, 50 mM KCl, 10 mM MgCl_2_) containing 1 μCi γ-^32^P-labelled ATP at 30 °C for 1 h, and then incubated with 1 to 2 μg MutT1 with or without 500 μM of 8-oxo-GDP, 8-oxo-GTP, GDP or GTP for 5,10 or 15 min at 30 °C. The reaction was terminated by adding 1× SDS-PAGE sample buffer (heating step was not included). The samples were resolved on 12% SDS-PAGE, and exposed to phosphor screen (Fujifilm Bas cassette2, Japan) for 4 h followed by imaging with Typhoon 9210 phosphor imager (GE Healthcare, USA) and Azure Sapphire (Azure Biosystems, USA).

### Phosphotransfer assay

Briefly, 1 μg of SK was incubated in kinase buffer (50 mM Tris-HCl, pH 8.0, 50 mM KCl, 10 mM MgCl_2_) containing 1 μCi of γ-^32^P-labelled ATP at 30 °C for 1 h, and then incubated with buffer, 2 μg *Msm*MutT1 WT or *Msm*MutT1 H170A for 1 h at 30 °C. The reactions were either terminated by adding 1× SDS-PAGE sample buffer (heating step was not included) or incubated with RR for 5 min at 30 °C for phosphotransfer activity, and then reactions were terminated as mentioned above. The samples were resolved on 12% SDS-PAGE. After electrophoresis, the gel was exposed to phosphor screen (Fujifilm Bas cassette2, Japan) for 4 h followed by imaging with Typhoon 9210 phosphor imager (GE Healthcare, USA) and Azure Sapphire (Azure Biosystems, USA).

### Quantification of the released ^32^*P_i_* from incubation of SenX-P with MutT1

Briefly, purified SenX3 (1 μg) was incubated in kinase buffer (50 mM Tris-HCl, pH 8.0, 50 mM KCl, 10 mM MgCl_2_) containing 1 μCi of γ-32P-ATP (3,000 Ci/mM) at 30°C for 2 h. Further incubation was with buffer, 2 μg MutT1 proteins for 1 h at 30°C. The reaction was stopped by 2% (final concentration) HCOOH. The reaction aliquots (2.5 μL) were spotted on PEI Cellulose F plate (MERCK # 105579), developed with 1.5 M KH_2_PO_4_ (pH 3.4), dried, exposed to phosphor imager screen (Fujifilm Bas cassette2, Japan) for 4 h, and imaged.

### Binding affinity with MST

The binding affinity of MutT1 with SKs was determined using microscale thermophoresis (MST) [8]. Briefly, 50 nM of phosphorylated or unphosphorylated forms of fluorescently tagged SKs namely, SenX3-mClover or PdtaS-GFP were titrated with MutT1 (concentration range mentioned in figure legends). The sample was then loaded into standard treated capillaries (Monolith^TM^ NT.115 series) and analyzed using a Monolith NT.115 (NanoTemper Technologies, Germany). The blue laser was used for a duration of 35s with excitation (MST power = 60%, LED power 40%). The data were analyzed using MO Control software (NanoTemper Technologies, Germany) to determine the K_D_ for the interacting proteins[8].

### Promoter activity assays

*M. smegmatis* strains were freshly electroporated with plasmids carrying the specific promoter constructs. For growth in acidic media, cells were first grown in 7H9 (pH 6.8) medium till OD_600_ of ∼0.6−0.8, harvested by centrifugation, resuspended in 7H9 (pH 4.7) medium, incubated at 37 °C for 3 h, harvested again and stored at −20 °C. For MOPS propionate media, the cells were grown in LBT media till OD_600_ of ∼0.6−0.8, transferred to the MOPS-propionate low phosphate media (final OD_600_ ∼0.2), incubated at 37 °C for 6 h, harvested and stored at - 20 °C [43]. The cells were resuspended in 500 ml 100 mM potassium phosphate buffer (pH 7.4), lysed by bead beating and centrifuged at 13K at 4 °C for 15 min. The total cell lysate proteins were quantified using Bradford assay. An aliquot of cell lysate (10 μg total protein) was incubated with 10 mM catechol in a 50 μl reaction, and incubated at 30 °C for 30 min. The absorbance at 375 nm was read in Tecan plate reader [44].

## Data Availability Statement

The data underlying this article will be shared on reasonable request to the corresponding author.

## Acknowledgements

The authors acknowledge laboratory colleagues for their critical comments on the manuscript.

## Credit authorship contributions statement

EAFE: Conceptualization, formal analysis, investigation, methodology, visualization, writing – original draft, writing – review & editing. UV: Conceptualization, formal analysis, methodology, supervision, funding acquisition, project administration, writing – review & editing. DPS and DKS: formal analysis, investigation, methodology, writing – review & editing.

---

**Fig. S1:**
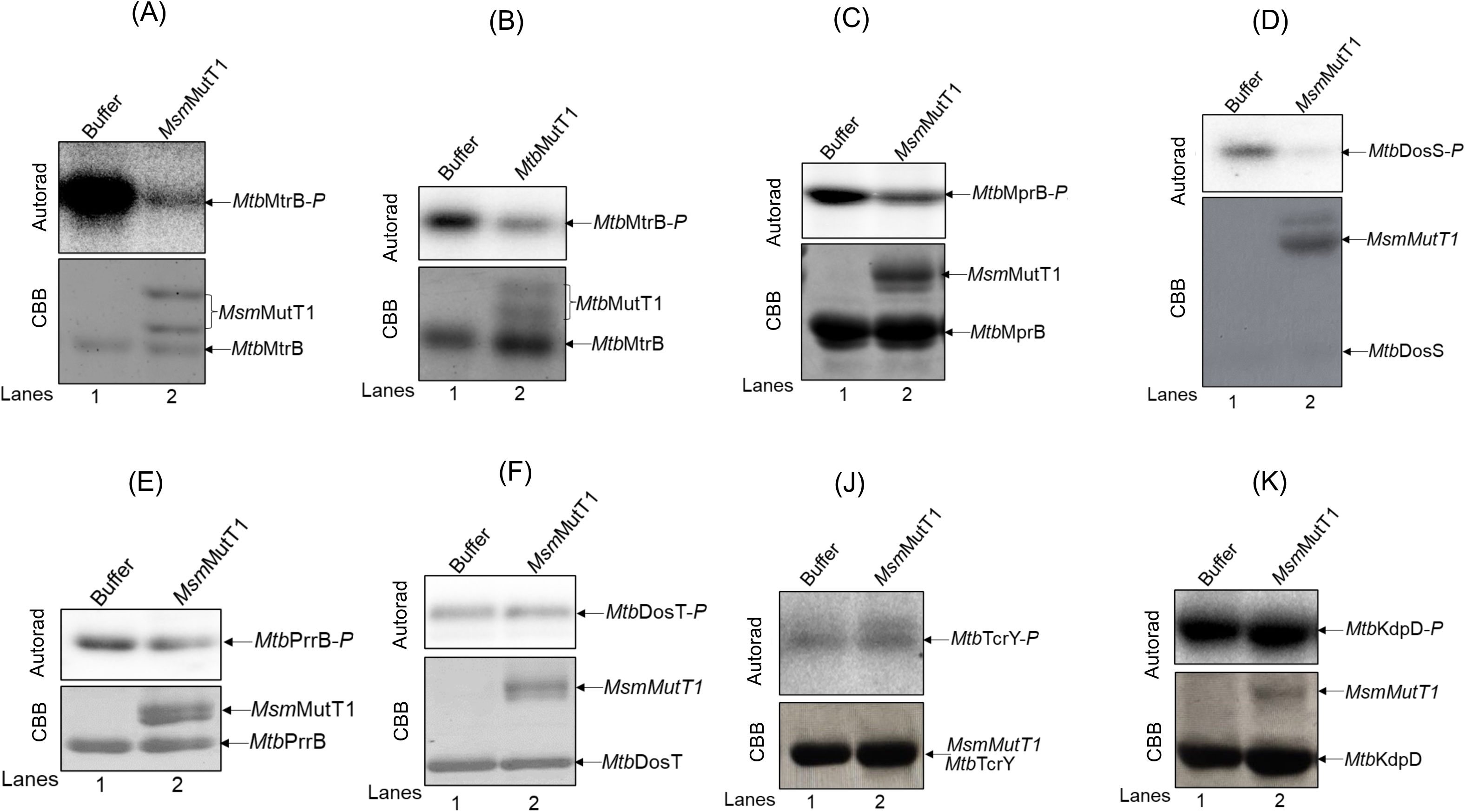
Identification of SKs which serves as a potential target for MutT1. Various phosphorylated SKs were incubated with *Msm* or *Mtb* MutT1 proteins, analyzed on SDS-PAGE, and autoradiographed (see methods section). (**A** and **B**) *Mtb*MtrB-*P* with *Msm*MutT1 (**A),** and *Mtb*MutT1 **(B**); (**C**) *Mtb*MprB-*P* with *Msm*MutT1. (**D**) *Mtb*DosS-*P* with *Msm*MutT1. (**E**) *Mtb*PrrB-*P* with *Msm*MutT1. (**F**) *Mtb*DosT-*P* with *Msm*MutT1. (**J**) *Mtb*TcrY-*P* with *Msm*MutT1. (**K**) *Mtb*KdpD-*P* with *Msm*MutT1. Top panels in A-K are autoradiograms and bottom panels are corresponding CBB stained gels.

**Fig. S2:**
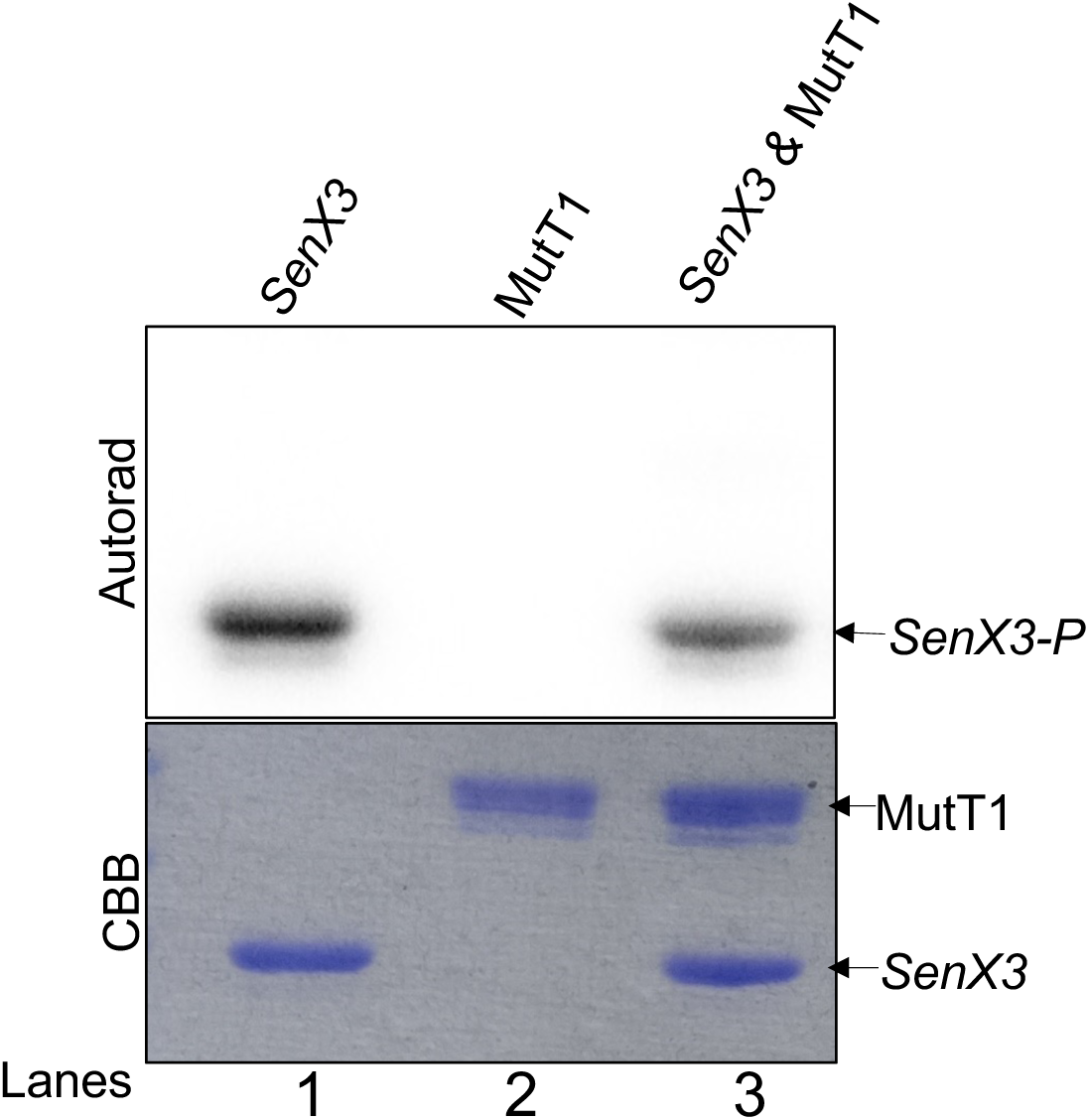
Analysis of MutT1 during SKs dephosphorylation process. For checking if MutT1 was phosphorylated while dephosphorylating SKs, the MutT1 proteins were incubated with either γ-^32^P-ATP or phosphorylated SK SenX3 and further incubated for 1 h at 30°C, analyzed on the SDS-PAGE and autoradiographed (see methods section). The top panel is autoradiogram and bottom panel is corresponding CBB stained gel.

**Fig. S3:**
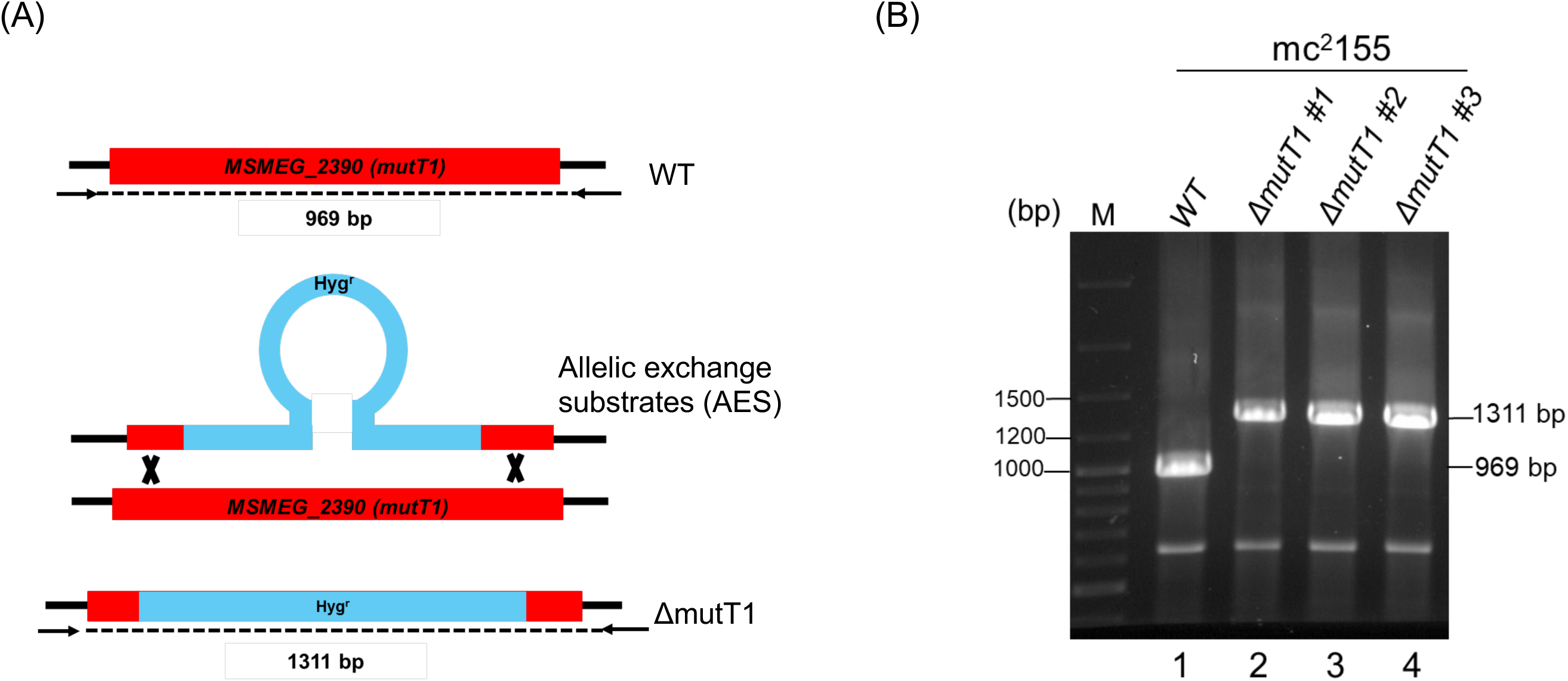
Generation of *mutT1* knockout in *M. smegmatis* mc^2^155 strain. **(A)** Schematic showing genomic organization of *MSMEG_2390* (*mutT1*) and Δ*2390* (*mutT1*)::Hyg loci with the binding sites of primers used for PCR based screening. **(B)** Representative agarose gel showing the amplification of *mutT1* locus, using the flanking primers, from the control mc^2^155, lane 1 and knockout strains (lanes 2, 3, and 4) where the wild type and *mutT1*::*hyg* alleles result in 969 bp and 1331 bp sized amplicons, respectively.

**Fig. S4:**
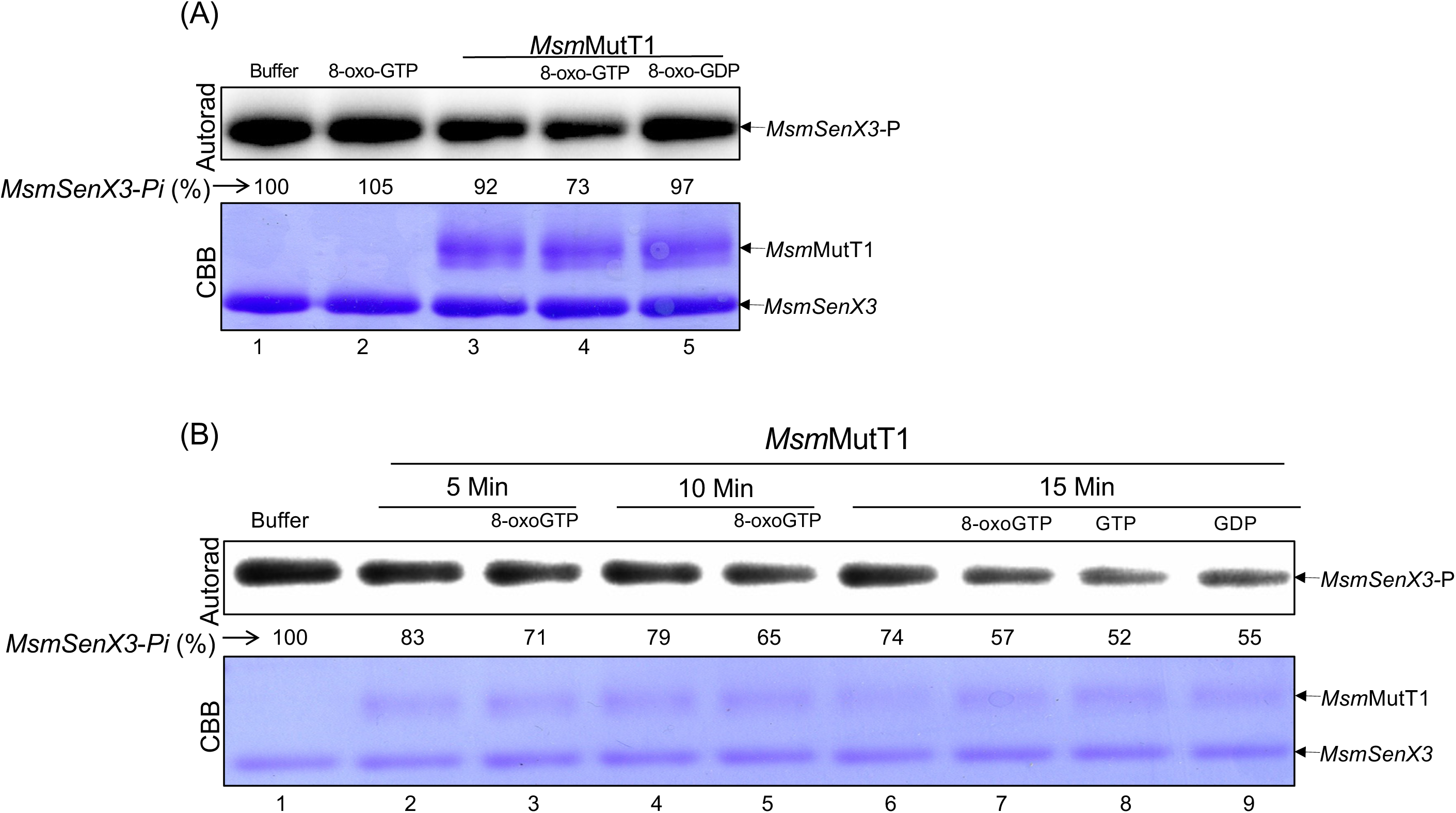
Role of MutT1 NTD binding substrates in regulating CTD mediated phosphatase activity: **(A)** The phosphorylated *Msm*SenX3 was incubated with either buffer alone, 500 μM 8-oxo-GTP or 8-oxo-GDP with or without *Msm*MutT1. **(B)** The phosphorylated *Msm*SenX3 was incubated with either buffer alone, 500 μM 8-oxo-GTP, GTP or GDP with or without *Msm*MutT1. The top panels in (A) and (B) are autoradiograms and bottom panels are corresponding CBB stained gels. The PAGE/Autoradiography images presented here is representative of three independent experiments.

**Fig. S5:**
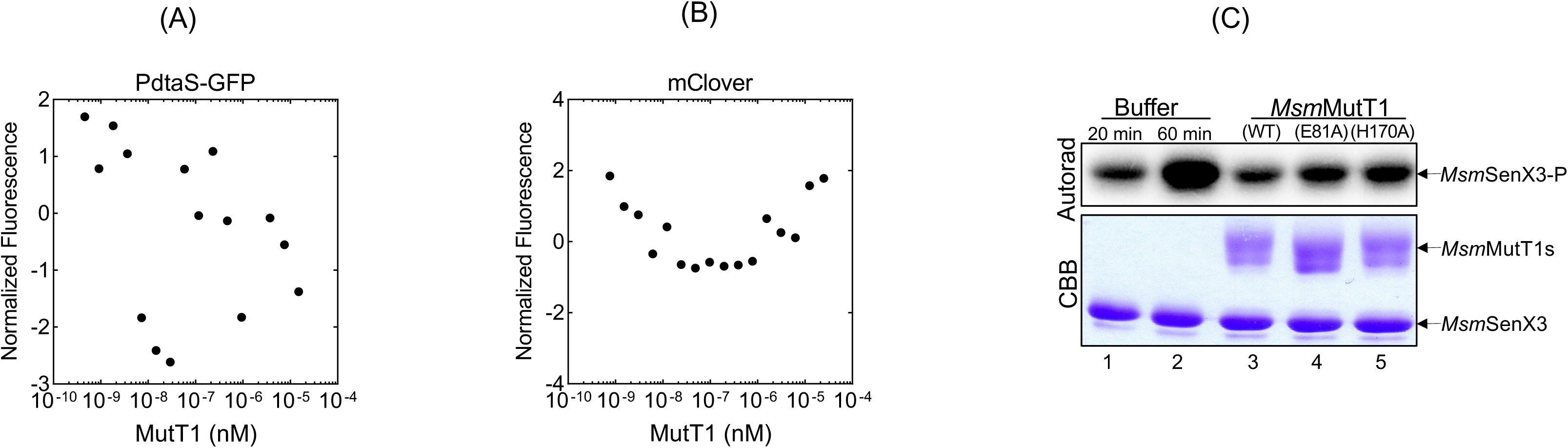
Interaction analysis of MutT1 with with non-substrate SK. **(A)** Normalized fluorescence intensity obtained from MST of 50 nM of PdtaS-GFP as a function of the concentration of the titrant MutT1 (concentration range- 0.45 nM to 15 μM). **(B)** Normalized fluorescence intensity obtained from MST of 50 nM of mClover as a function of the concentration of the titrant MutT1 (concentration range- 0.45 nM to 15 μM). **(C)** For checking the influence of the presence of MutT1 on autophosphorylation ability of SK, the *Msm*SenX3 was phosphorylated for 20 min at 30°C followed by either incubation with either buffer alone, *Msm*MutT1 (WT), *Msm*MutT1 (E81A), and *Msm*MutT1 (H170A) for additional 40 mins at 30°C and autoradiography was performed (see methods section). The top panel is autoradiogram and bottom panel is corresponding CBB stained gel.

**Fig. S6:**
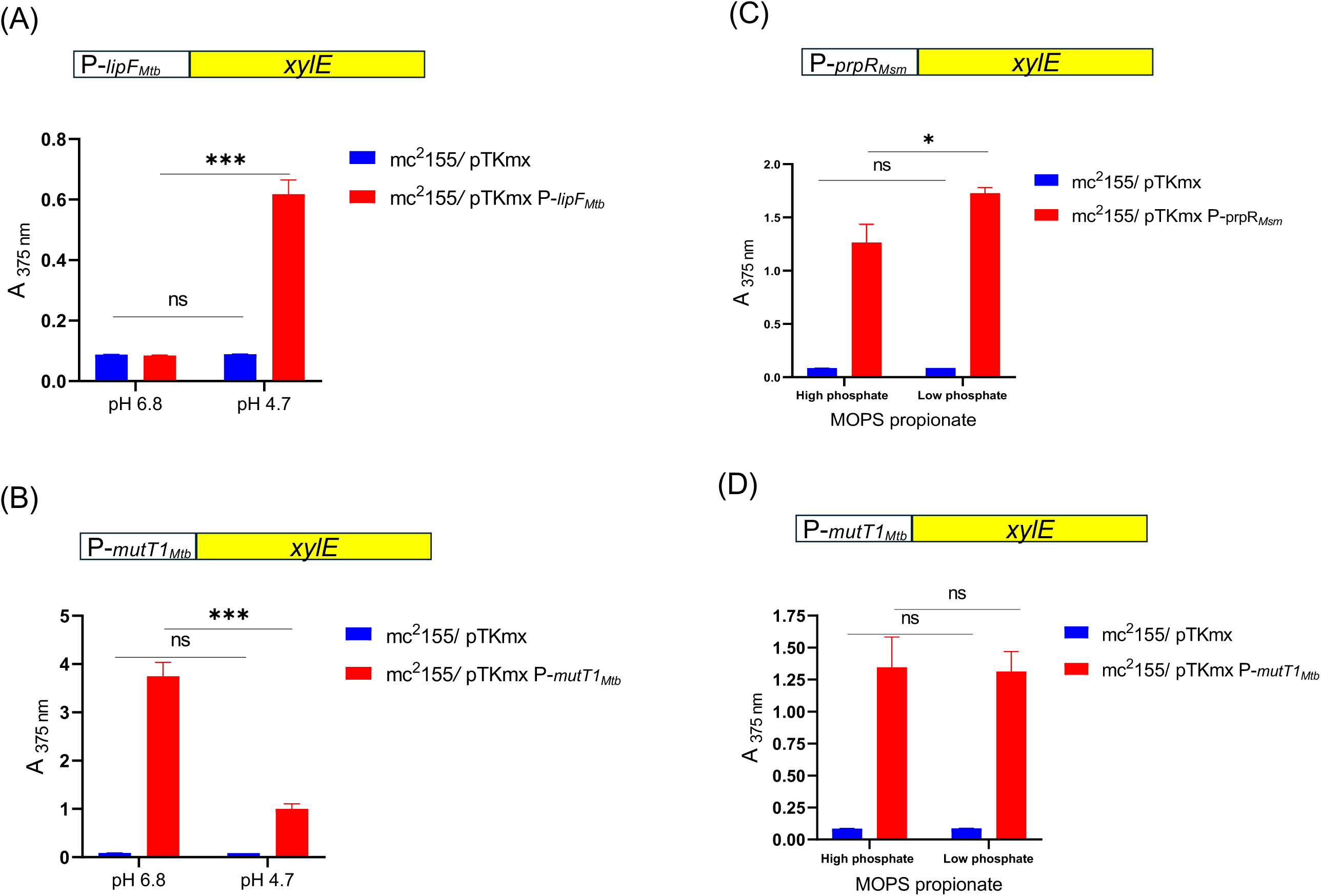
Investigation of the impact of MutT1 on TCSs target gene expression using *xylE* reporter. **(A)** *lipF* promoter was introduced upstream of the *xylE* ORF. The resulting reporter construct was electroporated in *Msm* mc^2^155/pTKmx (promoter less *xylE*) and *Msm* mc^2^155/pTKmx:P*_Mtb_*_lipF_ (*xylE* driven by *Mtb* lipF promoter). The cultures were grown in7H9 media of pH 6.8 or pH 4.7, and the cell-free extracts prepared and assayed for XylE activity. **(B)** *Mtb mutT1* promoter was introduced upstream of the *xylE* ORF and electroporated in *M. smegmatis*. *Msm* mc^2^155/pTKmx (promoter less *xylE*) and *Msm* mc^2^155/pTKmx: P*_Mtb_*_mutT1_ (*xylE* driven by *Mtb*mutT1 promoter) were grown in7H9 media of pH 6.8 or 4.7 and processed for XylE assay.. **(C)** *Msm prpR* promoter was introduced upstream of the *xylE* ORF and electroporated in *M. smegmatis*. *Msm* mc^2^155/pTKmx (promoter less *xylE*), and *Msm* mc^2^155/pTKmx:P*_Msm_*_prpR_ (*xylE* driven by *MsmprpR* promoter) were grown in MOPS propionate media with low phosphate (100 μM) or high phosphate (10 mM) and processed for XylE assay. **(D)** *Msm* mc^2^155/pTKmx and *Msm* mc^2^155/pTKmx:P*_Mtb_*_mutT1_ were grown in MOPS propionate media with low phosphate (100 μM) or high phosphate (10 mM) and processed for XylE assay. Bars represent mean ± SD for n = 3. One-way ANOVA method was used to calculate *p* value. *p* values, ∗ *p* < 0.05; ∗∗∗ *p* < 0.001 indicate significant differences between samples; ns- represent not significant.

**Fig. S7:**
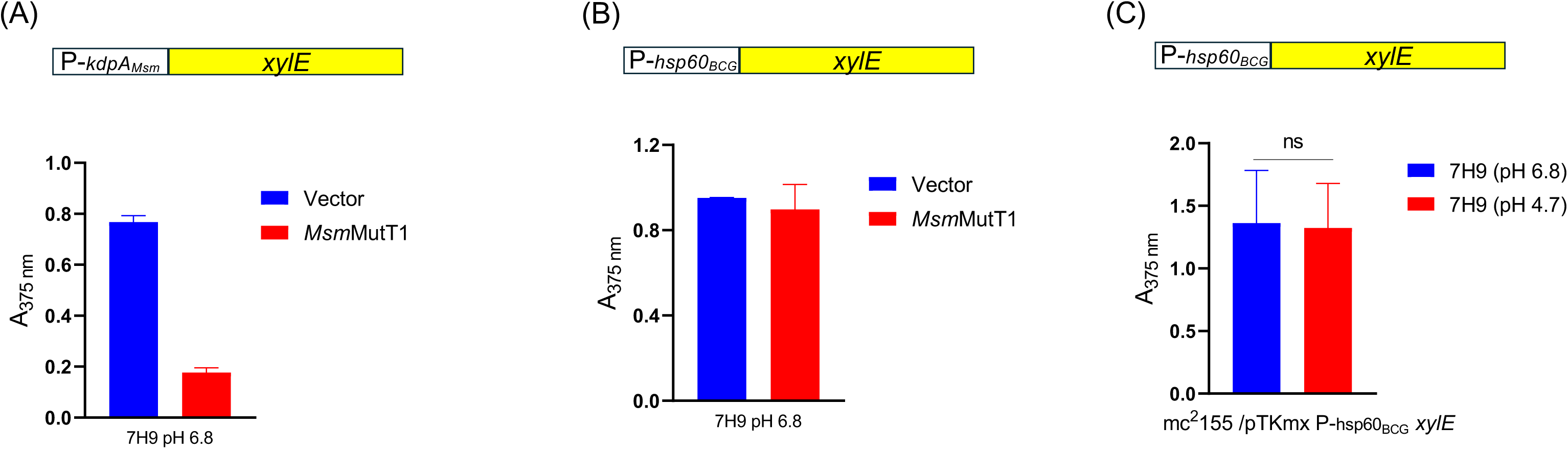
Examining specificity of MutT1 and its impact non-substrate SKs using *xylE* reporter. For this study, the *xylE* system was introduced under various promoters, **(A)** *Msm kdpA* promoter (a target of TCS KdpDE). **(B)** and **(C)** *M. bovis* BCG *hsp60* promoter. The generated reporter constructs were electroporated in the following strains, *Msm* mc^2^155 (WT)/ pCV125.1 and *Msm* mc^2^155 (WT))/ pCV125.1: *Msm*MutT1. The cultures were were grown in7H9 media at indicated pH- 6.8 or 4.7, and XylE activity was checked. The graph is representative of three independent experiments.

**Table S1:**
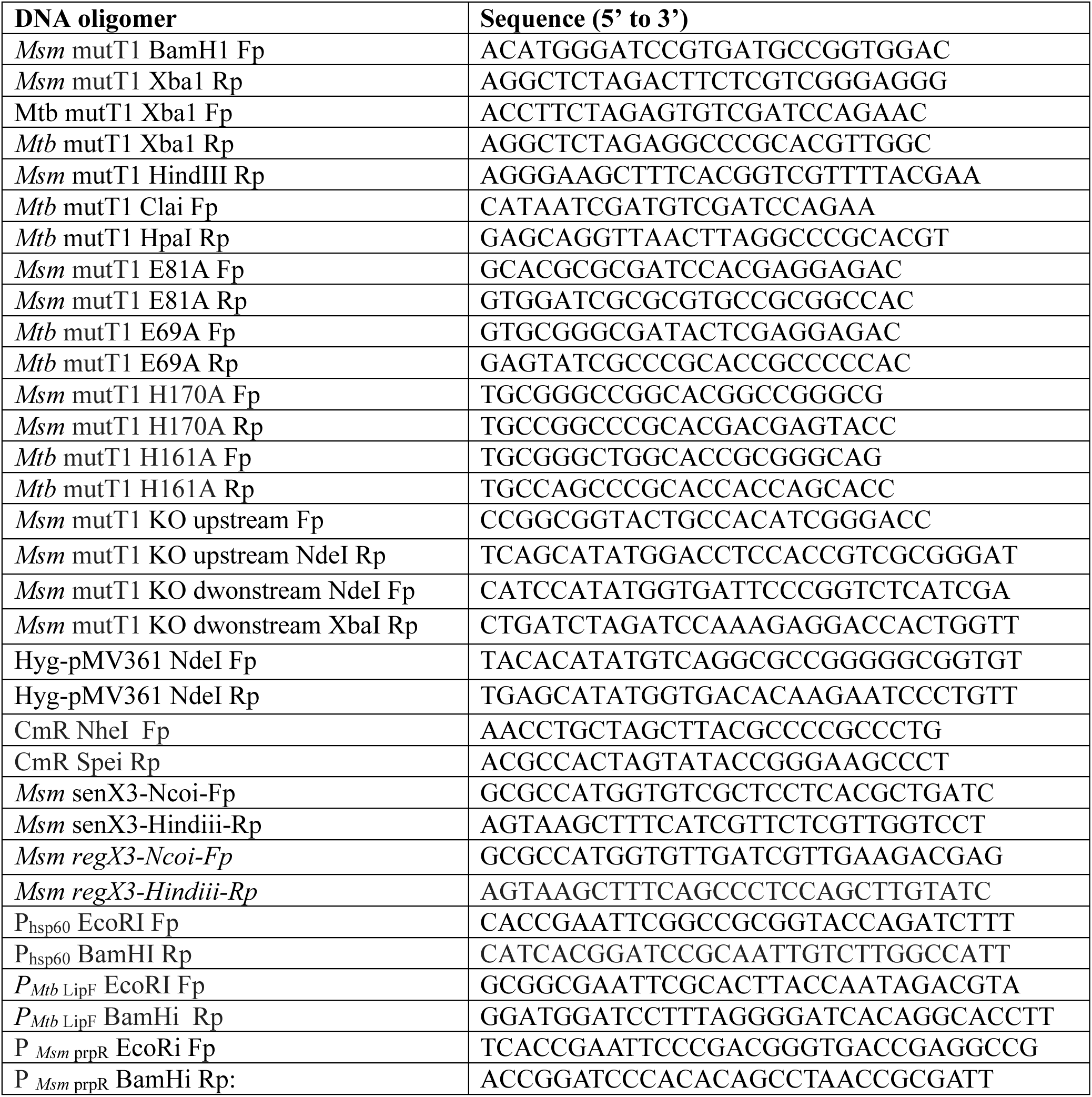
Description of oligonucleotides used in the study.

